# Floria: Fast and accurate strain haplotyping in metagenomes

**DOI:** 10.1101/2024.01.28.577669

**Authors:** Jim Shaw, Jean-Sebastien Gounot, Hanrong Chen, Niranjan Nagarajan, Yun William Yu

**Affiliations:** Department of Mathematics, University of Toronto; Genome Institute of Singapore, Agency for Science Technology, and Research (A*STAR); Yong Loo Lin School of Medicine, National University of Singapore; Computational Biology Department, Carnegie Mellon University

**Keywords:** Metagenomics, microbial strains, haplotyping, assembly

## Abstract

Shotgun metagenomics allows for direct analysis of microbial community genetics, but scalable computational methods for the recovery of bacterial strain genomes from microbiomes remains a key challenge. We introduce Floria, a novel method designed for rapid and accurate recovery of strain haplotypes from short and long-read metagenome sequencing data, based on minimum error correction (MEC) read clustering and a strain-preserving network flow model. Floria can function as a standalone haplotyping method, outputting alleles and reads that co-occur on the same strain, as well as an end-to-end read-to-assembly pipeline (Floria-PL) for strain-level assembly. Benchmarking evaluations on synthetic metagenomes showed that Floria is *>* 3*×* faster and recovers 21% more strain content than base-level assembly methods (Strainberry), while being over an order of magnitude faster when only phasing is required. Applying Floria to a set of 109 deeply sequenced nanopore metagenomes took *<*20 minutes on average per sample, and identified several species that have consistent strain heterogeneity. Applying Floria’s short-read haplotyping to a longitudinal gut metagenomics dataset revealed a dynamic multi-strain *Anaerostipes hadrus* community with frequent strain loss and emergence events over 636 days. With Floria, accurate haplotyping of metagenomic datasets takes mere minutes on standard workstations, paving the way for extensive strain-level metagenomic analyses.

**Availability:** Floria is available at https://github.com/bluenote-1577/floria, and the Floria-PL pipeline is available at https://github.com/jsgounot/Floria_analysis_workflow.

## 1. Introduction

To accurately assess the full genetic potential of microbial communities and capture their evolutionary and ecological dynamics, it is often necessary to resolve genomes at the strain level [1]. This is because there can be significant phenotypic variation between different strains of the same species [2,3], for example, *Escherichia coli* being either pathogenetic [4] or probiotic [5] in a strain-dependent manner. Thus, disambiguating strains within human microbiomes has important implications for human health and precision medicine [6,7]. Yet, many widely used metagenomic workflows [8,9,10] are not designed to recover multiple highly similar strain genomes. This often results in assemblies where less abundant strains are not recovered, potentially missing out on ecologically and medically important genetic features of the microbial diversity present.

Computational strain recovery comes in different forms. Strain *haplotyping* (or *phasing*) is the recovery of alleles, such as single nucleotide polymorphisms (SNPs), that co-occur along the same chromosome [11], or in the case of haploid microbes, the same strain in a community (haplotype) [12]. We are interested in using read overlap information to link alleles along the same strain haplotype. In this case, by clustering reads into strain-level clusters, one can also recover the sequence of alleles for each strain [13]. On the other hand, *strain-level assembly* is the recovery of *all* genomic content for each strain by base-level *de novo* assembly [14,15]. An orthogonal approach is *strain-level profiling*, where reference genomes from a database are used to identify corresponding strains from a metagenome [16,17,18]. Profiling assumes the existence of a reference genome for *each strain* in the population and does not reconstruct haplotypes explicitly. In this work, we will focus instead on phasing and assembly.

Short reads are limited in their ability to resolve between-strain similarities required to construct high-contiguity strain-level assemblies, but the advent of long-read Oxford Nanopore and PacBio sequencing has opened the door for more complete strain resolution. In particular, PacBio HiFi-based assembly methods leverage highly-accurate, but more expensive, long reads for strain-level assemblies [14,15], but HiFi reads are often not available for population-scale cohorts due to its cost. Therefore, methods that work well on long-reads with higher error rates, as well as short-reads, are still desirable despite their inherent limitations.

For nanopore reads, strain-level assembly is a challenging task at lower coverages or when using older, less accurate chemistries. Strain-level assembly is also computationally expensive for all technologies. In contrast, because phasing requires resolution of only a sparse subset of alleles, it is more feasible for sequencing data with lower accuracy, while being more computationally efficient than assembly-based approaches. Thus, phasing is a very useful task when strain-level assembly is too difficult or time-consuming. However, most existing haplotyping methods have only been designed and tested for short-reads [19,20,12,21,22]. A few long-read strain-level assemblers with built-in haplotyping capabilities exist, but they have yet to be applied to large cohorts with deep metagenome sequencing data [23,24].

In this work, we introduce *Floria*, a novel strain <u>haplotyping</u> algorithm that can take both long and short-read metagenomic datasets as input. With Floria, the haplotype phasing task takes only minutes on a standard workstation, enabling large-scale recovery of microbial haplotypes. Additionally, our package *Floria-PL* uses the Floria haplotyper to provide a one-command strain-level <u>assembly</u> pipeline for complex metagenomes that is more accurate and several times faster than existing methods.

## 2. Methods

Floria takes a set of mapped reads and single nucleotide polymorphisms (SNPs) and outputs a clustering of the reads into strain-level sets. Floria’s algorithm works by explicitly optimizing a haplotype phasing objective function in a local manner and then using a global network flow approach to retrieve strains (**Figure 1B**), relying on the fact that read coverage for a strain is consistent across the genome. Each resulting set of reads output by Floria (called a haploset) represents a group that is sequenced from the same strain. The consensus sequence of SNPs supported by a haploset (called a vartig) is also output, along with an estimate of the strain count for a species. The output haplosets can then optionally be assembled to obtain a phased, strain-level assembly. Pseudocode is available in **Supplementary Methods**.

Floria’s phasing algorithm is written in the Rust, a systems-level programming language, for speed. To facilitate Floria with all possible inputs, Floria has also been packaged into an end-to-end snakemake pipeline called Floria-PL (**Figure 1A**). Reads can be either sequentially assembled and mapped or analyzed via a classification approach to map reads to a sample-specific set of high-quality reference genomes. Mapped reads, whether to assembled contigs or reference genomes, are then used to call variants and processed by Floria’s core algorithm, which can be used to automatically generate phased assemblies (i.e., assembled haplosets) using different assemblers.

### 2.1. Preliminaries and MEC optimization

We first review the problem of polyploid haplotype phasing of a k-ploid organism [25], analogous to phasing *k* strains. We represent a read in variant-space as an element of the set {*−*, 0, 1, 2, 3}^*m*^ where *m* is the number of SNPs present in the contig. Let *r*[*i*] be the allele of the *i*th SNP in the read, where we set *r*[*i*] = *−* if the *i*th SNP is not covered by the read, *r*[*i*] = 0 for the reference allele, *r*[*i*] = 1 for the first alternate allele, and so forth. Paired-end reads are combined into one element in variant-space. We focus only on SNPs in Floria.

We represent the mapped reads as a set *R* = {*r*_1_, *r*_2_, …, *r*_*n*_}. One formulation of phasing is to partition *R* into *k* sets *R*_1_,, *R*_*k*_ where the set of reads *R*_*i*_ contains the reads coming from the *i*th haplotype, and this problem is commonly solved by optimizing the MEC model: given a partition of reads, the MEC score is the minimum number of alleles that must be changed in order to have an error-free phasing.

To define the MEC objective precisely, we first define *d*(*r*_*i*_, *r*_*j*_) as the number of non “*−*” alleles that differ between two reads and *s*(*r*_*i*_, *r*_*j*_) as the number of non “*−*” alleles that are the same between two reads. We define *H*(*R*_*i*_) to be the consensus haplotype of a set of reads: *H*(*R*_*i*_) is also an element in {*−*, 0, 1, 2, 3}^*m*^ where *H*(*R*_*i*_)[*j*] = *a* if the majority of reads have allele *a* at position *j*. If two alleles are equally supported, we can choose any of the majority alleles. We let *H*(*R*_*i*_)[*j*] = *−* if no read covers the jth allele (e.g., due to nucleotide deletions for a strain). Then given a partition *R*_1_,, *R*_*k*_ of *R*, the MEC score is

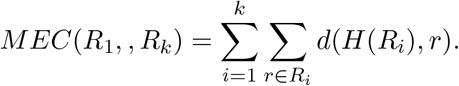

The MEC minimization problem is to find the partition that minimizes the MEC score; this turns out to be an NP-hard problem [26]. Thus, we solve it using a heuristic approach we describe below.

### 2.2. Local MEC optimization procedure through beam search

As described in **Figure 1B, step 1**, Floria first segments the contig into overlapping, contiguous regions of the genome. All reads that overlap each region is termed a *block*. Blocks are simply sets of reads, but *blocks are not necessarily disjoint*. The default block length, the length of the contiguous region corresponding to the block, is the 66th percentile of the read length, but it is adjustable by the user. The blocks overlap by 1/3 of the block length. In the subsequent sections, the read set *R* corresponds to all reads overlapping a specific block.

We first describe our method given a fixed number of strains, *k*. We optimize the MEC score based on the well-known heuristic beam search algorithm, variants of which have also been applied to polyploid haplotype phasing [27,28]. Our beam search routine is as follows: we start off by sorting the reads in *R* based on the first SNP the read covers and initialize a set of candidate solutions, *Solutions* = *P*, containing a single partition of k empty sets. We then iterate through every read *r*_*i*_ in order. During every iteration step, we create a new set of candidate partitions by adding *r*_*i*_ to every set in every partition in *Solutions*, giving us *k ·* |*Solutions*| candidate solutions. We then throw away identical partitions and keep the *γ* lowest MEC score solutions out of all *k ·* |*Solutions*| candidate solutions, where *γ* is some parameter. We repeat this procedure until we use all reads and then return the lowest MEC score partition in the final set of candidate solutions. By default, we keep *γ* = 10 candidates.

#### Optimizing the beam search

We introduce two additional heuristics beyond the basic beam search. Firstly, for the first 25 reads, we use *γ*^*′*^ = *k · γ* solutions instead to ensure a good initial phasing. Secondly, we use a new probabilistic heuristic to trim poor solutions. Let *D*_*ϵ*_(*r*_*i*_, *H*(*R*_*j*_)) be the exponentiated KL-divergence between the read *r*_*i*_ and the consensus haplotype *H*(*R*_*j*_) under an allele mismatch error probability *ϵ* as follows. First, let

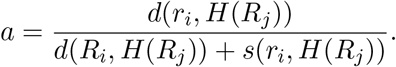

Here, *a* represents the fraction of differing alleles. Then, we define

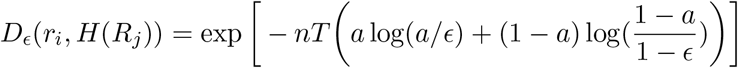

where *n* = *d*(*r*_*i*_, *H*(*R*_*j*_)) + *s*(*r*_*i*_, *H*(*R*_*j*_)) is the total number of SNPs shared by the haplotype and the read, and *T* is a constant that can be thought of as a “temperature” parameter that controls for the strictness of this divergence. By default, we set *T* = 0.25. *D*_*ϵ*_ will be small when the read *r*_*i*_ is dissimilar to the consensus haplotype, and can also be rigorously interpreted as a binomial test p-value in a limiting regime (Theorem 2 in [29]). Notably, the exponentiated divergence is minimized when *a* = *ϵ*, but we want it to be increasing when *a* is small, so Floria flips the sign when *a < ϵ* in the same manner as in a previous method [30].

We use *D*_*ϵ*_(*r*_*i*_, *H*(*R*_*j*_)) to prune unlikely sets to add a read to during beam search, thus tightening the search space to more plausible solutions. We define the relative goodness of the read assigned to set *R*_j_ in partition *P* as 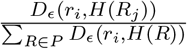. We then put *r*_*i*_ into the set *R*_*j*_ as a putative solution in our beam search only if the relative goodness is greater than a parameter which is 0.01.

Given the output of the beam search, we describe in Sections 1-3 in the **Supplementary Methods** additional heuristics for polishing the clusters and how to augment the MEC score with information from the *ϵ* parameter described in the previous section.

### 2.3. MEC Ratio as a threshold to detect local strain count

The above beam search algorithm outputs a partition for a fixed strain count and for a single block, but it requires prior knowledge of the number of strains present. To find the number of strains present automatically, we use an iterative heuristic that checks how the MEC score decreases as we increase a putative strain count. A simple probabilistic model of strain phasing (**Supplementary Methods 4**) was used to inform our heuristic method of finding the local strain count. Note that two strains may look almost identical in a local block. Therefore we will denote the number of strains that are distinct in this local block as the local strain count.

Let *MEC*^*∗*^(*k*) be the MEC score of the partition of the read set *R* into *k* sets as obtained by our MEC optimization procedure. Let *α*(*k*) : {1, 2, 3, …} *→* (0, 1) be a decreasing function of *k*, where we require (1 *− ϵ*)(1 + *α*(*k*)) *>* 1 for all *k*. The algorithm for determining the optimal strain count is as follows: we phase iteratively for increasing putative local strain count *k* = 2, 3… and check if 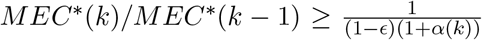 holds, i.e. the MEC score is still relatively large after increasing the putative strain count. If inequality holds, we stop phasing and return *k −* 1 as our true local strain count. We let *α*(*k*) = 1*/*[*k* + 1*/*3] by default and provide more options to the user depending on the desired resolution of phasing.

### 2.4. Flow graph construction on local haplotype blocks

We run the above iterative heuristic for all *N* blocks over the contig, obtaining a set of partitions *P*_1_, *P*_2_, …, *P*_*N*_ with low MEC scores. We then construct a directed acyclic graph (DAG) (**Figure 1B**, **step 3**) and a network flow capturing the flow of coverage between strains across blocks (**Figure 1B**, **step 4**). Let 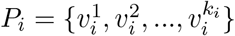 where 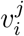 is the set of reads for the *j*th local strain in the *i*th block with local strain count *k*_*i*_. We let 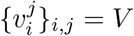 be the set of vertices in the graph and construct a directed edge between 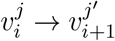 for all *i, j, j*^*′*^. We weigh the edge by the number of reads shared between 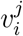 and 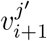 and remove the edge if the number of reads shared is less than 2.

This construction thus far does not describe a flow since the in-flow equals out-flow condition is not necessarily satisfied. We wish to find a reweighing of the edges that is faithful to the edge weights, yet is a true flow. Such a weighting gives a globally aware and thus robust signal that is not affected by local coverage variation or noisy reads being incorrectly partitioned (**Figure 1B**, **step 4**). This problem is similar to flow problems in transcriptome assembly, or viral quasispecies assembly [31,32,33], but we use a different objective function and path retrieval algorithm.

### 2.5. Exactly solving the flow problem as a linear program

Mathematically, given the graph as described in the previous section with *G* = (*V, E*) and weighting *w*, we wish to find a flow *f* on the edges that solves min_*f*_ ∑_*e*∈*E*_ |*w*(*e*) −*f* (*e*) | where *f* also has to satisfy the standard flow constraints: for all *ν* ∈ *V*, we require 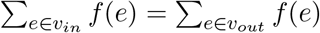with *ν* _*in*_ and *ν* _*out*_ being the edges into and out of a node.

This function is solvable by linear programming (LP). We first introduce auxiliary variables *t*(*e*) and the linear constraints *t*(*e*) *≥ f* (*e*) *− w*(*e*) and *t*(*e*) *≥ w*(*e*) *− f* (*e*). It follows that *t*(*e*) *≥* |*w*(*e*) *− f* (*e*)| and minimizing *t*(*e*) corresponds to minimizing |*w*(*e*) *− f* (*e*)|. Then we just solve the LP

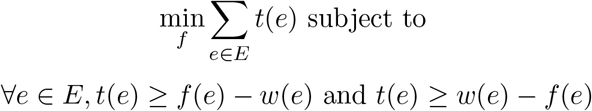

along with the original flow constraints on *f* to give us the resulting network flow. We solve this LP using the minilp [34] package, an open-source LP solver.

#### Conservative strain recovery by vertex-disjoint path decomposition

After obtaining a flow *f* on our DAG *G*, we iteratively extract vertex-disjoint paths from source to sink with the *largest minimum flow* (**Figure 1B**, **step 5**), removing each path from the graph after extraction. For each iteration, this is found in time *O*(|*V* ||*E*|) by a standard dynamic programming procedure along a DAG with traceback. We chose the largest minimum flow criterion because it is robust to coverage variation within the strains and high-coverage regions caused by mobile elements. Finally, for each path, we take the union of all reads within the path and output haplosets (**Figure 1B**, **step 6**). For each final haploset, we output the reads in the haploset and the consensus sequence of SNPs over all reads in the haploset, which we call the vartig. Since some reads may still be shared across haplosets, we assign each read to only its best haploset by finding the haploset, *Hap*, that minimizes *d*(*r, H*(*Hap*)).

Resulting phasings may be fragmented (**Figure 1B**, **step 5**), but due to the complexity of real metagenomes, we opted for a conservative *vertex-disjoint* path decomposition method as opposed to an *overlapping* path decomposition method [32] that is sometimes used for viral or transcriptome assembly. We initially tried using the standard greedy method of iteratively extracting paths with the largest flow and subtracting their flows to get an overlappping path decomposition. However, we found that we would get many duplicate vartigs due to overlapping paths forming from leakage of flow due to noise.

### 2.6. Estimating strain count and haplotype phasing quality (HAPQ)

For each contig phased by Floria, we output a statistic called the average strain count, or just strain count for short (not to be confused with local strain count). The strain count of a contig is the average number of haplosets covering each SNP. Unlike the number of strains in a community, strain count can be fractional, e.g. if two strains are identical for half of their genome but contain variation on the other half, the strain count is 1.5. Strain count thus represents a continuous estimate of the heterogeneity of the contig. We also output haplotype phasing qualities (HAPQ), analogous to mapping qualities, MAPQ, in read mapping. We describe our formula and interpretation of HAPQ in **Supplementary Methods 5**.

### 2.7. Floria-PL: a complete pipeline for haplotype phasing and assembly with Floria

Floria requires aligned reads and variants. To facilitate a phasing from reads directly, two approaches are implemented in the Floria-PL pipeline (**Figure 1A**): users can input a metagenomic assembly or reads can be classified against a Kraken database (UHGG in this work [35] v2.0.1) to retrieve highquality genome assemblies from species with an estimated coverage high enough for phasing (default 5X). We show the efficacy of this approach in **Supplementary Figure 1** and **9** for a spiked-in test and the synthetic dataset described below. Reads are mapped using minimap2 [36] against the selected reference genomes. For variant calling, both Longshot [37] (v0.4.1) and freebayes [38] (v2.11.0) are incorporated in Floria-PL. Additionally, haplosets output by Floria can be further processed to generate assemblies, and multiple assemblers were implemented within the pipeline: Megahit [39] (v1.2.9), Flye [40] (v2.7-b1585, kmer size = 16) and WTDBG2 [41] (v2.5). We further extended this approach to produce a comprehensive assessment pipeline to systematically compare phasing solutions (**Supplementary Figure 2**) covering read simulation, different mapping and variant calling tools, and a phased contigs assessment module.

**Fig. 1.**
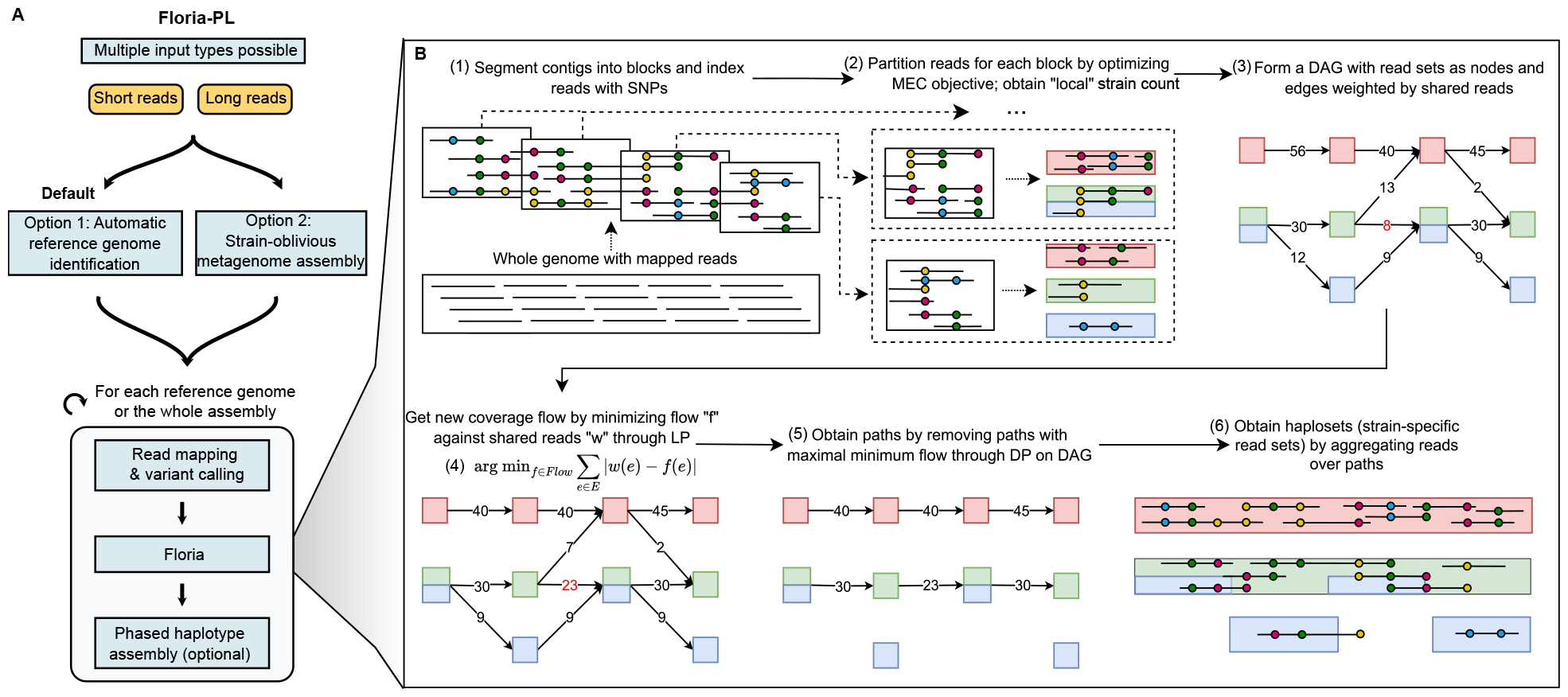
Overview of the phasing pipeline, Floria-PL, and core phasing algorithm, Floria. **A**. The Floria-PL pipeline first processes short or long-reads by either metagenomic assembly or by identifying a set of suitable reference genomes (default). Read mapping and variant calling are then performed. These outputs are then given to the Floria phasing algorithm. **B**. Floria optimizes a minimum error correction (MEC) model of SNP phasing locally and then finds a coverage-preserving network flow by linear programming (LP) on a directed acyclic graph (DAG) constructed from the locally phased blocks. Floria outputs strain-level read sets called haplosets and their haplotypes, called vartigs, for downstream analysis. Haplosets can be assembled to give haplotigs (phased contigs).

**Fig. 2.**
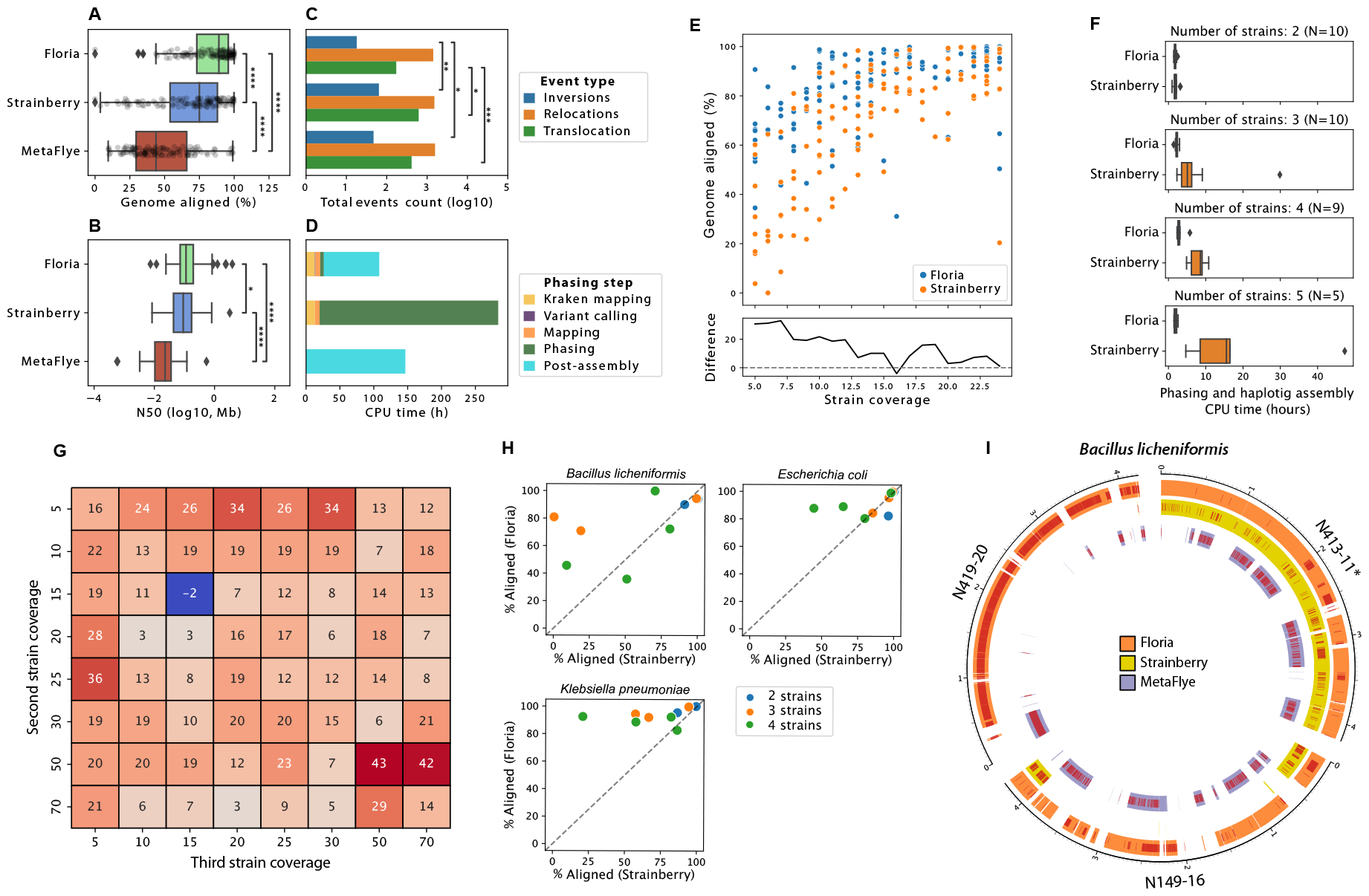
Assessment of phased assemblies for long-read synthetic communities and real reads. **A**. Percent aligned (i.e., covered by assemblies) for all strain genomes, **B**. N50 (log10), **C**. number of structural variants within the strain assemblies, and **D**. CPU time spent with Floria, Strainberry and MetaFlye for a nanopore synthetic read community of 40 species with 120 strains. **E**. Percent aligned of all 120 strain genomes as a function of strain coverage. The lower graph displays the average difference between the two methods. **F**. Phasing and assembly runtime as a function of the number of strains for each species. **G**. Phased assembly improvement (absolute difference) based on percentage aligned for Floria relative to Strainberry for a synthetic community with 3 *K. pneumonia* strains. The first strain has 15× coverage. **H**. Phased assembly results for pooled, real nanopore reads from multiple strains, showing the aligned percentage of the assembled strains. **I**. Circos plot [42] of assemblies obtained from a synthetic 3-strain community of pooled, real nanopore reads for *B. licheniformis*. Bands represent segments of the true genomes with alignments from the assemblies, and red lines are SNPs. N413-11 (marked with an asterisk) was used as the reference genome for mapping and phasing.

### 2.8. Synthetic communities generation and performance benchmarking

To produce a realistic strain synthetic community, we selected the 40 most abundant species in a cohort of 109 Singapore gut metagenome samples [43], based on the median abundance obtained from Kraken classifications against the UHGG database. For each species, only genomes with completeness *>* 90% and contamination lower than *<* 5% were kept and sorted using the score: Completeness - 5 × Contamination + 0.05 × log(N50). The 500 strains with the highest score for each species were retained and compared using skani [44] (v0.1.0). The resulting pairwise ANI distances were processed to generate strain clusters using an agglomerative clustering (scikit-learn v1.2, 1% ANI threshold, average linkage), and genomes with the highest score within their cluster were assigned as the strain cluster representative. To explore a large panel of strain numbers, the synthetic community was generated by putting together differing numbers of strains for different species, stratified into 5 groups (number of species / number of strains): 5/1, 10/2, 10/3, 10/4, and 5/5. Genome coverage was defined randomly with a minimal coverage of 5X and a maximal coverage of 25X using a discrete uniform distribution. Nanopore reads were simulated with badread’s [45] nanopore2020 model at 87.5% mean identity. Description of other synthetic communities can be found in **Supplementary methods 7**.

Phased assembly results generated from samples with known isolates were assessed in a similar fashion to Strainberry’s methodology [23]. Briefly, query assemblies were compared against each reference strain’s genomes using MUMmer4 [46], and each contig was assigned to the closest isolate using a coverage × identity score. Only alignments covering at least 50% of their query were considered. Aligned contigs were concatenated and compared all together against their assigned true genome with MUMmer4 to produce GAGE [47] assembly evaluation metrics, including genome coverage and identity, percent aligned, duplication ratio, and potential structural variants. We primarily benchmarked against Strainberry and metaFlye. Strainy [24] is another promising recently developed long-read phasing tool, but it is designed for phasing “one or a few bacterial species” currently and extending to larger metagenomes is a “work in progress” [48], so we did not benchmark against it in its current version. Benchmarking of CPU and memory usage is further described in **Supplementary methods 8**.

## 3. Results

### 3.1. Floria improves recovery and runtime for long-read phased assembly of diverse strain mixtures

We first compared Floria to MetaFlye and Strainberry on a 40 gut species (120 strains) synthetic community (see **Methods**), using the automatic reference genome identification approach (**Figure 1A**, option 1) and Flye as Floria’s downstream assembler. Floria improves phased assembly results (**Figure 2A, Supplementary Figure 3**), with a significantly larger portion of strain genomes recovered (21% relative mean improvement, Mann-Whitney U Test, p-value *<*1×10-6, **Supplementary Figure 4A**), higher contiguity compared to Strainberry (**Figure 2B**, **Supplementary Figure 4B**) and higher completeness, with an assembly size on average 23% larger (mean = 3.4Mb and 2.7Mb for Floria and Stainberry respectively). Assemblies are not only larger but also have improved quality, with a lower number of misassemblies, especially for translocations (mean = 1.4 vs 5.2 for Floria and Strainberry respectively, **Figure 2C**). Floria’s phasing is fast, with only 5.3 CPU hours being used to phase samples, and an additional 82 CPU hours to assemble haplosets with Flye, in comparison to 263 CPU hours with Strainberry (phasing and assembly) and 146 CPU hours with MetaFlye (**Figure 2D**). Any assembler can be used downstream for Floria depending on the user’s preference. Using WTDBG2 instead of Flye, we observed a significant reduction of duplication ratio and misassemblies at the cost of genome completeness, though genome completeness still remained higher than Strainberry (**Supplementary Figure 3**). We found that Floria is more suitable for phasing and assembling strains at lower coverage compared to Strainberry, with up to a 33% relative increase in strain genome recovery at 7× coverage (**Figure 2E**). The difference in performance is more pronounced when a species has many strains, with Floria being able to phase and assemble species with 3 or more strains much more completely (+19% absolute difference of strain genome aligned on average, Mann-Whitney U test p-value *<*3×10-8, **Supplementary Figure 5**).

**Fig. 3.**
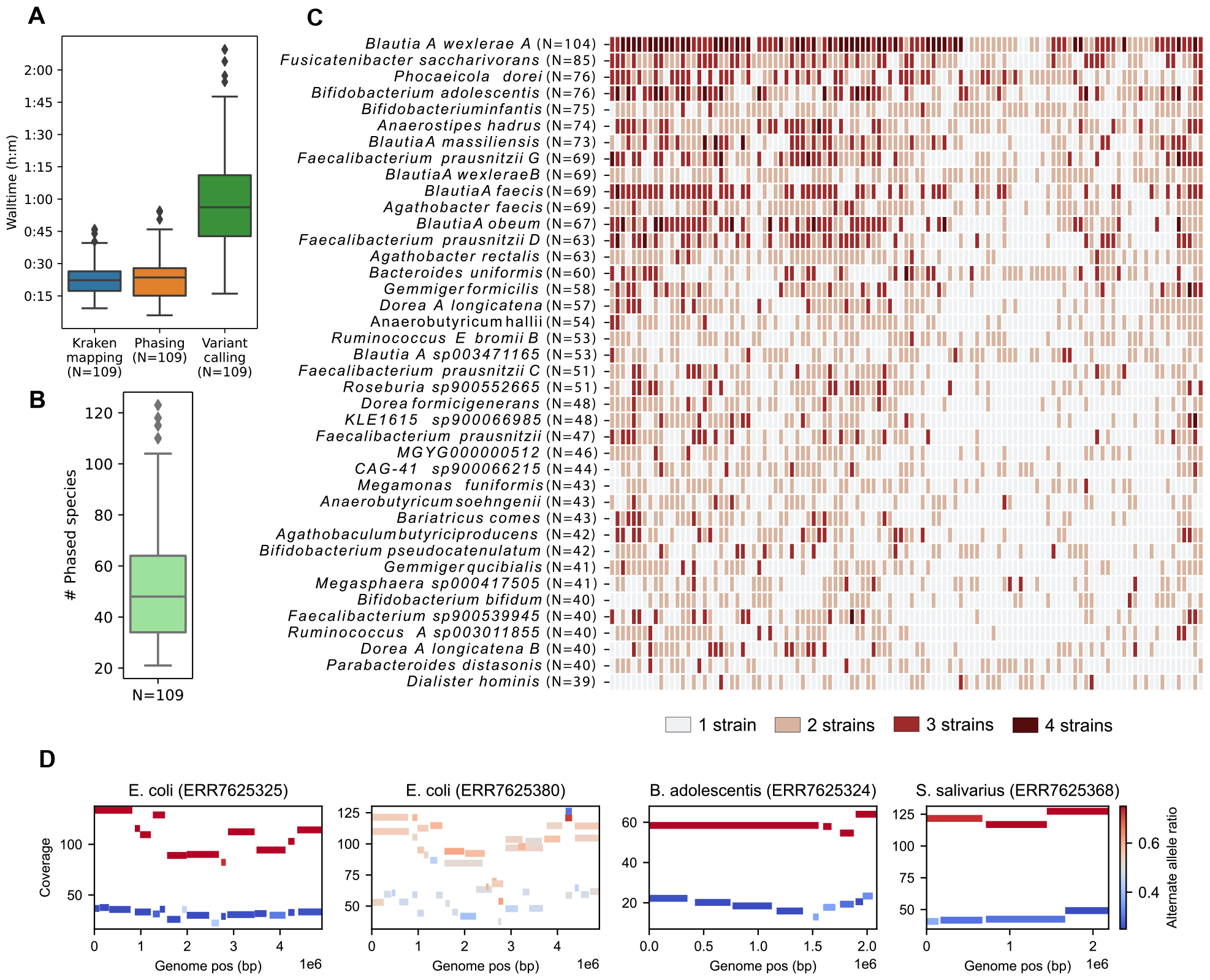
Phasing of 109 gut nanopore metagenomes. **A**. Runtime of each major component of the phasing pipeline for each sample. **B**. Number of species that have been phased by Floria for all 109 SPMP samples. **C**. Heatmap of the number of strains within the population for the 40 most abundant species based on short-read abundance. Species are sorted by prevalence (i.e. the # of samples a species appears). **D**. Coverage-vartig plots of Floria phasings with sample and species denoted. Each bar is a vartig that spans positions shown in the x-axis and is colored by its fraction of alternate alleles.

**Fig. 4.**
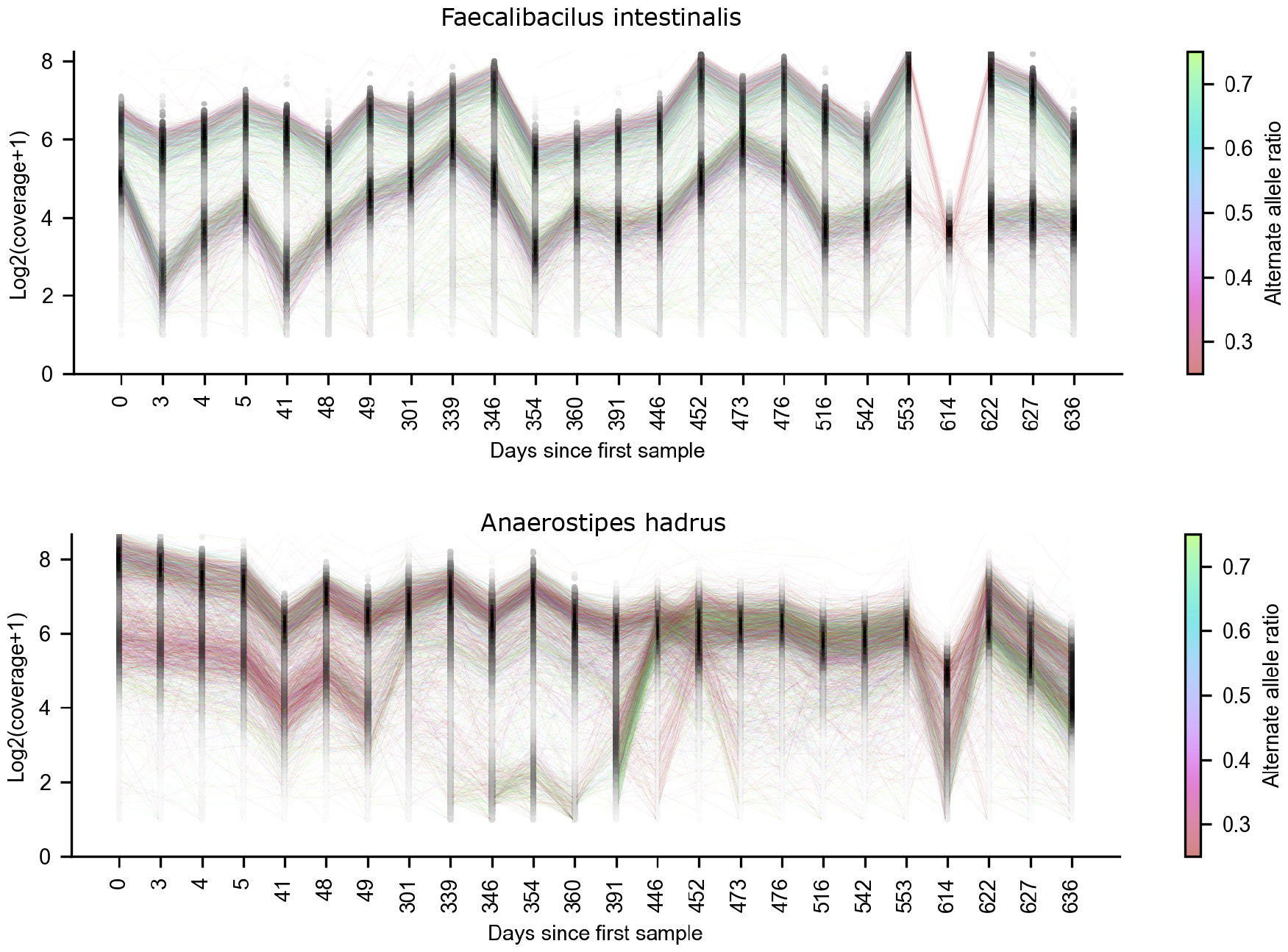
Strain tracking of longitudinal samples from Donor A in Watson et al [51] for *Faecalibacilus intestinalis* and *Anaerostipes hadrus* strains. Each line represents a vartig-vartig correspondence colored by the percentage of alternate alleles relative to the reference for the paired vartigs. The density of vartigs is shown in black along each sampling date. The coverage is log2(x+1) transformed to stabilize variance [52] Note the non-uniformity of the sampling dates.

We found that Strainberry’s runtime also increases substantially as the number of strains increases, while Floria’s runtime remains low (**Figure 2F**). Applying Floria directly on a MetaFlye assembly (**Figure 1A**, option 2), we observed a considerable improvement of the strain genomes recovery from the initial assembly (34% relative mean improvement, **Supplementary Figure 3**), but with a decreased performance compared to the reference genomes based phasing. This approach remains useful for metagenomic samples without a comprehensive reference genome database available and is currently limited to Floria, as Strainberry is unable to directly phase a metagenomic assembly with a mixture of species of different strain counts.

To investigate the impact of coverage on a sample, we simulated reads for three *Klebsiella pneumonia* strains at variable coverage. Floria recovers more strain content on average (89.6% vs 74.5% for Floria and Strainberry respectively) in almost all coverage configurations (**Figure 2G**) while Strainberry can not assemble low-abundance strains when variation in coverage is high (**Supplementary Figure 6**). Using 7 *E. coli* strains with increasing divergence, Floria recovers a higher proportion of strain genomes, even for closely related strains compared to Strainberry (**Supplementary Figure 7**). Next, we constructed synthetic communities using real reads from multiple isolates from 3 species: *Bacillus licheniformis* (PRJNA1029794), *Klebsiella pneumoniae* (PRJNA1033449), and *E. coli* [49]. For each isolate, we first assembled high-quality ground truths for quality assessment. For all species, Floria showed improvement in genome completeness over Strainberry (**Figure 2H**), with most of the strains being resolved at more than 80% completeness (mean of 88% and 75% genome aligned for Floria and Strainberry, respectively). In some cases, Floria is able to reconstruct almost the entire strain genome with relatively good accuracy (average ANI=99.5%) and a low number of misassemblies (average=4.6 structural variants), while MetaFlye and Strainberry generated either one strain or an incomplete mixture of all strains (**Figure 2I, Supplementary Figure 8**).

### 3.2. Exploring diversity with nanopore sequencing of 109 gut metagenomes using Floria

To explore Floria’s utility for studying strain diversity within diverse microbial communities, we analyzed data from a population study of 109 gut microbiomes with deep nanopore sequencing [43]. On average, Floria-PL took *<*2 hours per sample (*<*25 minutes for Floria phasing) using 8 CPUs (**Figure 3A**). We analyzed *>*5,000 sample-species pairs with sufficient coverage (*>*5× by default, **Supplementary Figure 9**), yielding phasings for *>*50 species on average per sample (**Figure 3B**). The strain number within a species-sample pair has been defined using Floria’s strain count estimate with haplotype quality (HAPQ) superior or equal to 15, subsequently length-normalized across all contigs within a genome and rounded to an integer (**Supplementary Methods 6, Figure 3C**). The ability to detect multiple strains for a species in the gut microbiome was correlated with the prevalence (the number of samples the species appears in) of the corresponding genome (**Supplementary Figure 10D**). Correspondingly, more abundant species such as *Blautia wexelerae A, Bifidobacterium adolescentis*, and *Faecalibacterium prausnitzii G* tend to have a higher proportion of samples with multiple strains (*>*50%).

We visualize four phasings with high coverage: a 2-strain and a 3-strain *E. coli* community, a 2-strain *B. adolescentis* community, and a 2-strain *Streptococcus salivarius* community (**Figure 3D**). For the three 2-strain communities, differential strain coverage confirms our phasings; these haplosets can in principle be binned to give near-complete haplosets. We also inspected the 3-strain *E. coli* community with the IGV [50] (**Supplementary Figure 11**) and found concordant haplosets for three strains. This shows Floria’s potential as a method of obtaining phasings and quickly confirming them without any assembly.

### 3.3. Short-read phasing enables tracking of multi-strain dynamics in longitudinal samples

We describe additional benchmarks showing that Floria can also phase short-reads in **Supplementary Methods 9** and **Supplementary Figure 12**. Leveraging short-read phasing capabilities, we investigated a longitudinal set of 24 human gut short-read sequencing samples collected over 636 days from a healthy individual labeled “Donor A” from Watson et al [51]. For these samples, we called variants with freebayes (with the `--pooled-continuous` option) on the combined BAM file and phased each sample on the combined VCF file. We then tracked the vartigs across samples with a reciprocal mapping procedure (**Supplementary Methods 10**). Floria took less than two hours (20 threads) and 11 GB of RAM to haplotype all 24 samples.

We focus on two species with multiple strains and high abundance across all samples, *Faecalibacilus intestinalis* and *Anaerostipes hadrus* (**Figure 4**). For *Faecalibacillus intestinalis*, a recently isolated species [53] from the human gut with limited characterization, exactly one major strain and one minor strain were present across 636 days. To confirm that there were indeed two strains present, we visualized the haplosets output by Floria for the first time point in IGV (**Supplementary Figure 13**). The remarkable stability observed here could reflect distinct niches for these strains [54], but other explanations are possible as stability and competition have a nuanced relationship [55].

As an example of a species with lower stability and interesting gain/loss patterns, we tracked the dynamics of *Anaerostipes hadrus*, a butyrate-producer with potential benefits for intestinal health, but butyrate-producer phenotypes are highly strain-dependent [56]. This analysis highlighted the emergence of a low abundance strain that transitioned to become a high abundance strain between days 391 and 446. To confirm this emergence, we visualized Floria’s outputs during day 391 and day 446 (**Supplementary Figure 14**) in IGV, showing that low-coverage haplosets on day 391 have identical alleles as the high-coverage haplosets on day 446. This emergent strain was not the same as the minor strain on day 0, as indicated by the high number of alternate alleles in the emergent strain, whereas the minor strain on day 0 contains mostly reference alleles (**Figure 4**). There are thus at least 3 strains present over this timescale. Notably, the low-coverage strain on day 391 has *<*1/15th of the coverage of the major strain, demonstrating that even low-abundance strains can be detected and tracked with Floria. Finally, we observed that for both species, strains observed at other time points were lost only in the sample for day 614. This could be an indication of a substantial perturbation to the gut microbiome at the strain level, but more likely could be an indication of mislabeling for the day 614 sample.

## 4. Conclusion

The importance of strain-level analyses, combined with ever-increasing amounts of sequencing data, implies that accurate and efficient methods for unraveling strains will be of continued interest for the foreseeable future. In response, we developed Floria, a metagenome haplotype phasing tool, along with an associated phasing pipeline that can rapidly extract microbial haplotypes for downstream analysis. Floria’s phasing and assembly capabilities provide a path forward for large-scale analyses of diverse metagenomic data with even short and noisy long-reads.

## Acknowledgments

This work was supported by the A*STAR Computational Resource Centre through the use of its high performance computing facilities and the Natural Sciences and Engineering Research Council of Canada (NSERC) grant RGPIN-2022-03074. J.S was supported by an NSERC CGS-D scholarship.

