## Supplementary Materials for "Floria: Fast and accurate strain haplotyping in metagenomes"

**Algorithm 1:** Floria phasing algorithm pseudocode

**Input** : BAM, Contig, VCF
**Output:** Haplosets $Hap_1, ..., Hap_\ell$ for Contig

$\epsilon \leftarrow$ Provided by user or estimated(BAM)
$\mathcal{R} = \{R, ...\} \leftarrow$ Break contig into chunks and let each $R \in \mathcal{R}$ be the set of reads mapped to a chunk
Partitions = {}
### parallel for loop below
**for** $R$ *in* $\mathcal{R}$ **do**
  $MEC^* = []$; one-indexed array
  $Parts_R = []$; one-indexed array
  **for** $i = 1$ *to MaxPloidy* **do**
    $P \leftarrow$ BeamSearch($R$,$\epsilon$, $i$)
    $P \leftarrow$ IterativeOptimize($P$,$\epsilon$)
    $MEC^*[i] \leftarrow$ Get$\epsilon-$MECScore($P$, $\epsilon$)
    $Parts_R[i] \leftarrow P$
    **if** $MEC^*(i) <$ *NumberOfAlleles(R)* $\cdot\epsilon$ **then**
      $BestPart \leftarrow P$
      Break
    **end**
    $alpha(k) = 1/(k + 1/3)$
    **if** $MEC^*(i)/MEC^*(i-1) \geq \frac{1}{(1-\epsilon)(1+\alpha(i))}$ *and* $i \geq 2$ **then**
      $BestPart \leftarrow Parts_R[i-1]$
      Break
    **end**
    $BestPart \leftarrow P$
  **end**
  Partitions $\leftarrow$ Partitions $\cup\{BestPart\}$
**end**
$(V, E) \leftarrow$ Construct Weighted DAG By Shared Reads(Partitions)
$w \leftarrow$ Construct Flow Via Linear Program$(V, E)$
$Hap_1, \ldots, Hap_\ell$ = Get Paths With Max Min Flow Via DP$(V, E, w)$
Return $Hap_1, \ldots, Hap_\ell$

#### 1 MEC score degeneracy and the epsilon-MEC score

In the beam search procedure, the initial starting partition is empty. This is an issue for MEC optimization because putting a read in an empty set is always the optimal greedy move as it does not increase the MEC score. We would prefer having similar reads cluster together even if one of the sets is empty. To combat this degeneracy, we define the epsilon-MEC score. Let $\epsilon$-MEC(R1,, Rk) = MEC(R1,,Rk) + S where S is the number of consensus alleles that only have a single read supporting it and is the previously defined error parameter. The epsilon-MEC score regularizes the degeneracy issue, as putting a read into an empty set increases the score by the expected number of errors in the read. We thus replace the MEC score in the beam phasing procedure with the epsilon-MEC score, knowing that the $\epsilon$-MEC score instead of MEC implicitly.

#### 2 Final iterative optimization of partitions

Given the output of the beam phasing procedure, we further refine the partition by a Kernighan-Lin like algorithm [30]. For each read in each set of the partition, we check if moving the read to another set decreases the total epsilon-MEC score. If it does, we store this move and the change in score. After checking all (k-1)—R— possible moves, we execute either 1/10 of the best scoring moves or 1/3 of the best scoring moves if there are ¡ 10 possible moves. We repeat this 10 times or until the epsilon-MEC score does not improve anymore.

### 3 Automatic parameter estimation of epsilon

Given that epsilon is an important parameter for both the epsilon-MEC score and beam search based phasing, we provide a heuristic to automatically estimate epsilon from data. From the input BAM file supplied by the user, we look at the pileup table for an arbitrary contig and obtain a distribution of the mismatch error rate. We then take the 66th percentile of the mismatch error rate as epsilon, which is slightly higher than the median estimate so as to account for reference bias and miscalled SNPs. We use this estimate at the start of Floria, and use it for all subsequent calculations. The user can also provide a specified epsilon value if prior information is known for their sequencing technology.

### 4 MEC score ploidy detection

Below we give a heuristic derivation of our local ploidy detection algorithm, which is based on a simple probabilistic model with assumptions on the MEC score. While the use the $\epsilon$-MEC score for phasing during the beam-search procedure, now we use the standard MEC score.

Suppose that we have reads from $k$-strains $R_1, ..., R_k$ and are trying to partition the $k-$strains into $n \leq k$ sets. We ask: what is the expected MEC score in this case?

We need some assumptions to make this tractable. Let $C_i$ be the average coverage of each strain (i.e. average coverage of each base in the strain) and let $C = \sum_i C_i$ with $C_1 > C_2 > C_3 ....$ We'll also assume that $C_i > \sum_{j>i} C_i$, i.e. the strain abundance decreases quickly. While this assumption is not true in general, for the special case of two strains, it is almost always true.

Let's assume that the algorithm works by initially putting $R_1, ..., R_n$, the $n$ most abundant strains, into separate sets in the partition and then lumping $R_{n+1}, ..., R_k$ into the set which minimizes the MEC. Let us assume that the reads call alleles with error rate $\epsilon$. Then the expected MEC score is

$$\mathbb{E}[MEC(n,k)] \approx \sum_{i=1}^{n} \epsilon * C_i * L + \sum_{j=n+1}^{k} (1-\epsilon) * C_j * \min_i d(H(R_j), H(R_i)).$$

The first $n$ terms are just allelic errors due to read errors. The second sum comes from lumping $H_j$ into the set with $H_i$ such that the distance between the two haplotypes is minimized; recall that $d(H(R_j), H(R_i))$ is the number of differing alleles between the two strains. The $1-\epsilon$ is the probability an allele is correct, in this case meaning there is an error because the two haplotypes differ. There are also possible errors from the $n+1, ..., k$ strains actually having the same allele as strain $i$, but the alleles are miscalled – we ignore these errors and assume this is relatively small. Note that we use the assumption $C_n > \sum_{j>n} C_j$ because implicit is the fact that even lumping all low abundance strains together does not change the consensus allele.

#### 4.1 Ratio of MEC scores

We can use the above formula to answer the question: assume that we have $n$ strains. What is the expected change in MEC score from an $n-1$ partition to an $n-$partition? The above formula gives:

$$\frac{\mathbb{E}[MEC(n,n)]}{\mathbb{E}[MEC(n-1,n)]} = \frac{\epsilon C L}{\sum_{i=1}^{n-1} \epsilon * C_i + (1-\epsilon) * C_n * \min_i d(H_n, H_i)}.$$

Rewriting this and letting $\min_i d(H_n, H_i) = d_n$, we get

$$= \frac{1}{(C - C_n)/C + (1-\epsilon) C_n/(C\epsilon) * (d_n/L)}.$$

Now we note don't want to detect strains that have too low coverage. We want the smallest covered strain to have at least $\epsilon C$ coverage otherwise it is indistinguishable from errors. Thus we let $C_n \geq \epsilon C$. Plugging in this inequality yields

$$\geq \frac{1}{(1-\epsilon)(1 + d_n/L)}. \quad (1)$$

Thus equation 1 serves as a lower bound for what this ratio should be. Making an assumption that this upper bound $\geq$ is actually a reasonable approximation $\approx$, we can choose this value as a sort of decision boundary.

Intuitively, if the real MEC score has a smaller ratio than this decision boundary, the MEC score has decreased more than we expected, meaning that the new phasing is very good. If the MEC score has not decreased as much as we expected, maybe there are only $n-1$ strains in the phasing.

##### 4.2 Algorithm

A simple algorithm to determine ploidy is to simply check $MEC(n,n)/MEC(n-1,n)$, the MEC scores from the beam-search phasing and see if it has a greater ratio than $\frac{1}{(1-\epsilon)(1+d_n/L)}$. If the real output ratio is greater, then we assume the ploidy is actually $n-1$, since it doesn't decrease as much as we expect. If the real output ratio is smaller than this conservative estimate, then we assume that $n$ is a better estimate of the number of strains than $n-1$.

##### 4.3 Empirical value for $d_n/L$

The value $d_n/L$ represents the fraction of alleles that differ. Denote $d_n/L = \alpha_n$. While $\alpha_n$ depends on the exact strains considered, we can set it to an empirical function. Importantly, it should be decreasing as a function of $n$, because it is a minimum over $n$ values. In floria v0.0.1, we have three possible functions for $\alpha_n$ corresponding to the option `--ploidy-sensitivity = 1,2,3`. The options 1,2,3 correspond to $\alpha_n = \frac{1}{n^a+b}$ where $a = 0.5, 1, 1$ and $b = 1, 1/3, 1$, so $\alpha_n$ is decreasing, which makes the above ratio larger – so the algorithm attempts to phase more strains. By default, we set the sensitivity to level 2.

#### 5 HAPQ score

Spurious haplosets arising due to errors in our automatic clustering may occur. We seek to quantify haploset goodness by a measure we call HAPQ, analogous to MAPQ for read alignment. Like MAPQ, HAPQ is a measure of how likely it is that the haplotype arises due to error and should be filtered out.

Suppose we have haplosets $Hap_1$ and $Hap_2$, which are sets of reads. Let $C(Hap_1, Hap_2)$ be the interval containment of $Hap_1$ in $Hap_2$, meaning that if $Hap_1$ spans SNPs in $[a,b]$, and $Hap_2$ spans SNPs in $[c,d]$,

$$C(Hap_1, Hap_2) = \frac{|[a,b] \cap [c,d]|}{b-a+1}$$

where $a, b, c, d$ are SNP indices along the contig.

Let $s(Hap_1, Hap_2) = s(H(Hap_1), H(Hap_2))$ be the number of identical alleles shared between the two consensus haplotypes of the haplosets (i.e., the vartigs). Let $L$ be the number of SNPs in the block, and $[a,b]$ defined as before for $Hap_1$. Then

$$\text{HAPQ}(Hap_1) = \min(60, t_1 * t_2 * t_3)$$

where

$$\begin{aligned} t_1 &= \max_{Hap_i, i\neq 1} 40 \cdot \left[ (1 - C(Hap_1, Hap_i)) \frac{s(Hap_1, Hap_i)}{s(Hap_1, Hap_i) + d(Hap1, Hapi)} \right], \\ t_2 &= \min(1, |Hap_1|/3), \\ t_3 &= \ln((b-a)/L + 1). \end{aligned}$$

The term $t_1$ gives a low HAPQ if two haplosets look very similar to another haploset and one is highly contained in the other, meaning that this haploset should maybe be a part of another haploset instead. $t_2$ makes sure there are enough reads in the haploset, and $t_3$ gives a high HAPQ if the vartig is long. Our formula is heuristic but is inspired by minimap2's formula for MAPQ [1], which is also a product of three terms with similar interpretations.

#### 6 Floria strains count estimation

Assuming $n$ contigs from a species, with size $c$ and associated Floria's strain count $s$ of minimum 1, we can define a species-level strain count (SSN) within a sample as the contigs size-adjusted mean of the strain counts:

$$\text{SNN}(Spe_1) = \frac{\sum_{i=1}^{n} (\max s_i, 1) * c_i}{\sum_{i=1}^{n} (c_i)}$$

To validate this estimator for different HAPQ values, we calculated the species strain counts for each species within the 40-species synthetic community and compared the value to the real strain numbers (**Supplementary Figure 15**). The estimation with $HAPQ > 15$ displays the most accurate result (Spearman correlation coefficient = 0.92), with a slight underestimation of the number of species for species with 2 or more strains, increasing with the number of strains. To avoid an overestimation of the strain count within real samples, we used a similar HAPQ threshold for the SPMP strain count estimation. For the SPMP dataset analysis, SNN values for each species have been rounded to the nearest integer to obtain an approximated but more practical representation of the number of strains for each sample-species pair.

### 7 Simpler communities design

Other synthetic communities were produced as follows. The 3 *Klebsiella pneumonia* community is composed of ASM1990034v1 (1st), ASM1990032v1 (2nd) and ASM1990030v1 (3rd). Nanopore reads were simulated in a similar fashion than before, 1st strain coverage has been fixed to 15X. The 7 *E. coli* strains community is similar to Strainberry dataset, with 20X coverage for K-12 strain (used as reference) and 30X for others. For both communities, Flye has been used for the post-phasing assembly as described before.

For real reads communities, subgroups of size 2, 3 and 4 strains were constructed using the following strains order: *E. coli*: [`'786605'`, `'542093'`, `'623214'`, `'899091'`], *B. licheniformis*: [`'N419_barcode16'`, `'N413_barcode03'`, `'N413_barcode11'`, `'N419_barcode20'`], *K. pneumonia*: [`'N319_barcode04'`, `'N319_barcode12'`, `'N325_barcode01'`, `'N326_barcode18'`]. For less than 4 strains, strains used are picked from left to right (e.g: `'786605'`, `'542093'`). Strain coverage follows the same principle based on the coverage gradient [25, 20, 15, 30]. Reads were assembled by Flye (v. 2.9.2-b1786) with the nano-raw input types and default options. Assemblies were further polished with Medaka (v. 1.7.2) consensus method and model `'r941_min_hac_g507'`. Reads were provided to the subsampling pipeline available with Floria subsampling pipeline for automatic reads subsampling (seed = 50) and phasing with same software versions used for the synthetic metagenome analysis. Haplotigs were assembled using the Flye assembler.

### 8 Computational resources benchmarking

Benchmarking of all phasing software and major components of the phasing processes was based on snakemake benchmarking module results. We decided here to use a CPU normalized wall time metric instead of the directly provided CPU time metric for benchmarking tool performance. This decision was made based on three factors: First, phased assembly solutions tested here rely on multiple steps with inconsistent CPU usages, leading to intermittent and sometimes very long but low CPU loading (i.e, a period when the defined number of CPUs is underused) such as for Strainberry's variant calling step. Second, such pipelines are more and more run on cloud computing units for which one of the main factors of consideration is the instance lifespan. Last, we assume here that the instances system CPU usage is negligible compared to the high CPU usage of most of the phased assembly software. All phased assembly results generated during the phasing assessment steps were done on a c6a12x AWS instance with the exception of StrainXPress for which a r5a12x was used to accommodate its high memory usage. Most of the multi-threaded steps were done with 48 CPUs, except when references were split and for which only 8 CPUs were used for each step. SPMP phasing has been processed on a separate cluster powered by Intel Xeon Platinum 8268 48-Cores 2.90 GHz processors with 192GB DDR4 2933MHz available.

### 9 Floria short-read phasing benchmark

While Floria produces long, accurate haplotypes for long-read data, many metagenomic samples only have short-read sequences available. Leveraging the same 120-strain synthetic community as before (section **Synthetic metagenome generation**), using ART[2] and we assessed Floria's short-read phasing and assembly against StrainXPress, a strain-level assembler, and the strain-oblivious assembler Megahit.

Floria assembles more strains (mean 72% aligned, Supplementary Figure 12) than Megahit (mean 50% aligned) but less than StrainXPress (mean 84% aligned). Nevertheless, this improvement for StrainXPress comes at the cost of a fragmented assembly with more than 6.9×105 sequences (against 3.5×105 and 1×105 for Floria and Megahit respectively) and a higher duplication ratio (1.32 and 1.18 for StrainXPress and Floria respectively). Furthermore, Floria requires drastically fewer computational resources than StrainXPress. Floria is more than 6 times faster than StrainXPress when considering variant calling, mapping, and haplotyping, and is more than 2 times faster when including base-level assembly after haplotyping (Supplementary Figure 12). StrainXPress takes more than 243 GB of memory compared to 29 GB for Floria, severely limiting its usage outside large computer clusters.

### 10 Reciprocal vartig mapping algorithm

For a pair of samples, we deduced corresponding vartigs between the two samples by the following reciprocal mapping procedure. First, for one of the samples which we call the "query" sample, we scan over each vartig in the query and find vartigs in the other "reference" sample for which this query vartig shares $> 2$ SNPs and has no errors – we call this a **hit**, giving us a set of hits for each query vartig. Second, we find a maximal sequence of non-overlapping hits subject to the following score:

$$Score(V_i, V_j) = s(V_i, V_j) - 0.2 * d(V_i, V_j).$$

Here $V_i, V_j$ are the vartigs on the SNP-level, and the $s, d$ functions are the same as before. For each query vartig, we get a sequence of hits with their respective scores. To get the maximal sequence of non-overlapping hits, we can use a standard dynamic programming. This returns a sequence of optimal hits for the query vartig to a set of reference vartigs. To get the final correspondence, we swap the query and reference and redo the procedure. We take these optimal hits that are present in both the original and swapped outputs to be our final correspondences.

Once we had our optimal reciprocal hits, we only retained hits with 0 errors, or with $< 95\%$ error rate (i.e. 1 differing SNP for every 20 SNPs) and used these hits to define strain-level vartig correspondences.

Our vartig aligner is available in the following repository: `https://github.com/bluenote-1577/vartig-utils`.

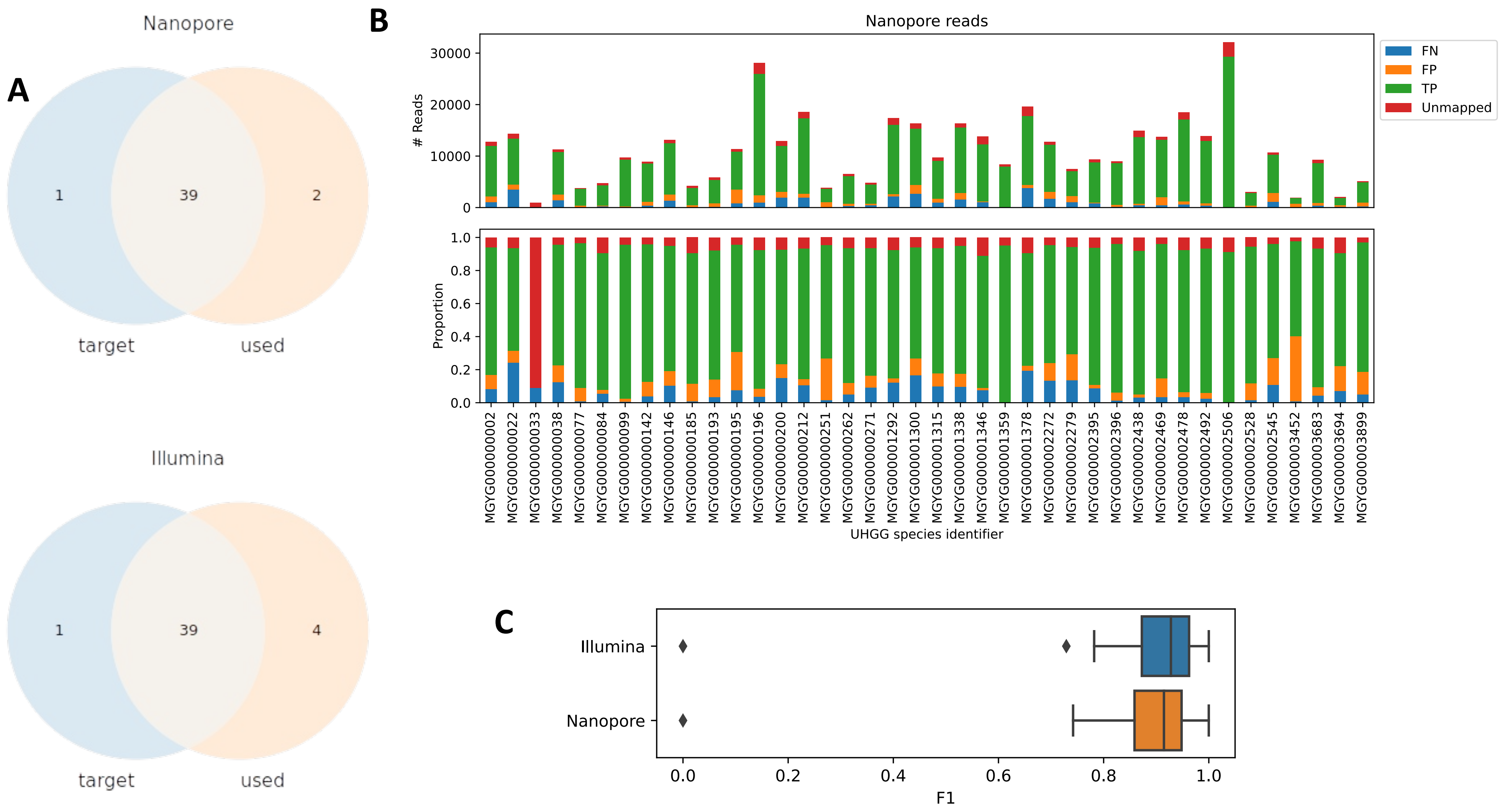


**Supplementary Figure 1. Kraken mapping procedure on 40 species, 120 strain metagenome.** A. Venn diagram representing the intersection between the species that have been used for the 40 species, 120 strain synthetic metagenome (target) and species detected by the kraken procedure (used). B. Number of reads and their proportion for each reference genome. FP = mapped read comes from another species. TP = mapped read comes from the same species. FN = the species' read is mapped to an another genome. C. Boxplot of the resulting F1 scores obtained the above read mapping results.

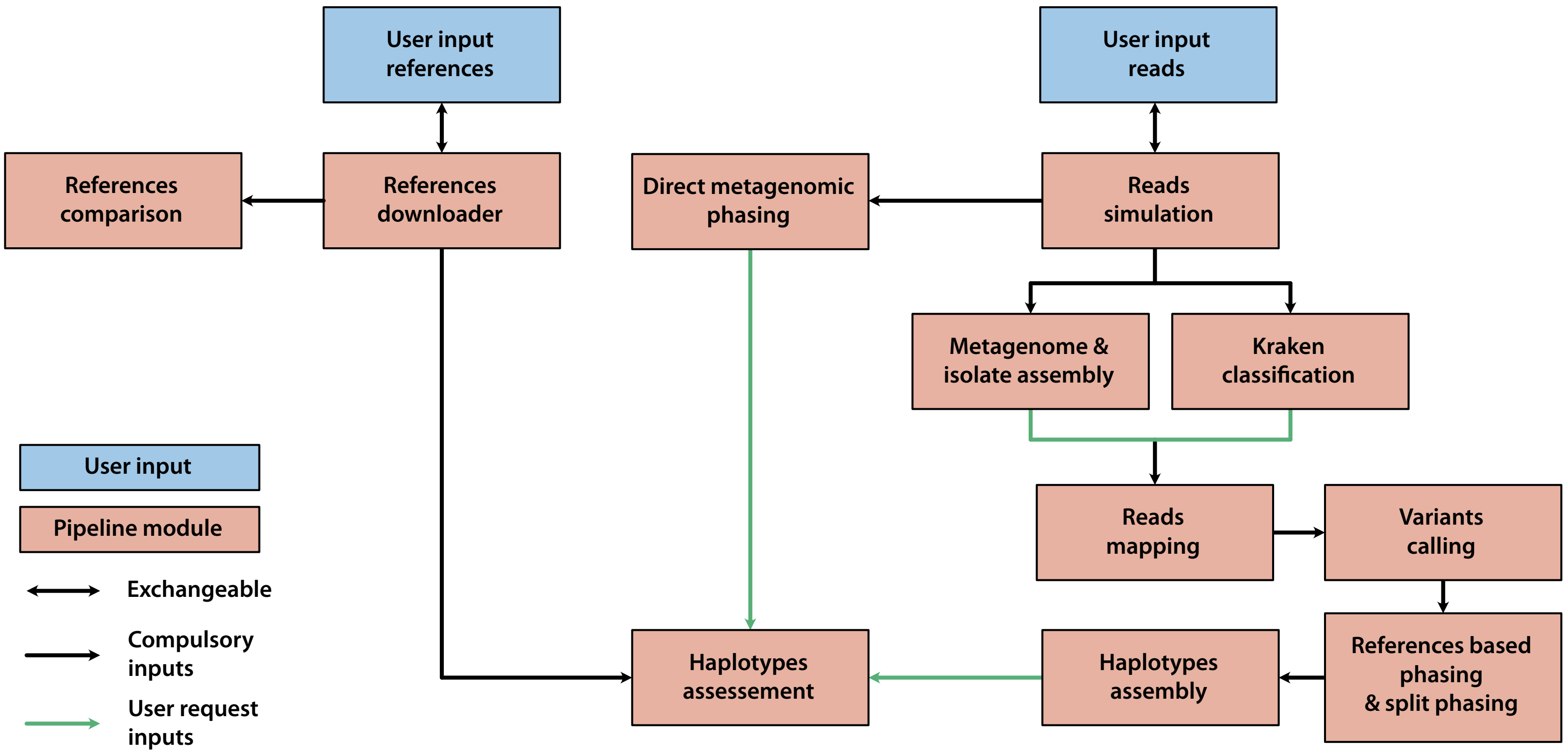


**Supplementary Figure 2. Schematic representation of the assessment pipeline.** Each orange block represents a module and blue blocks are user inputs.

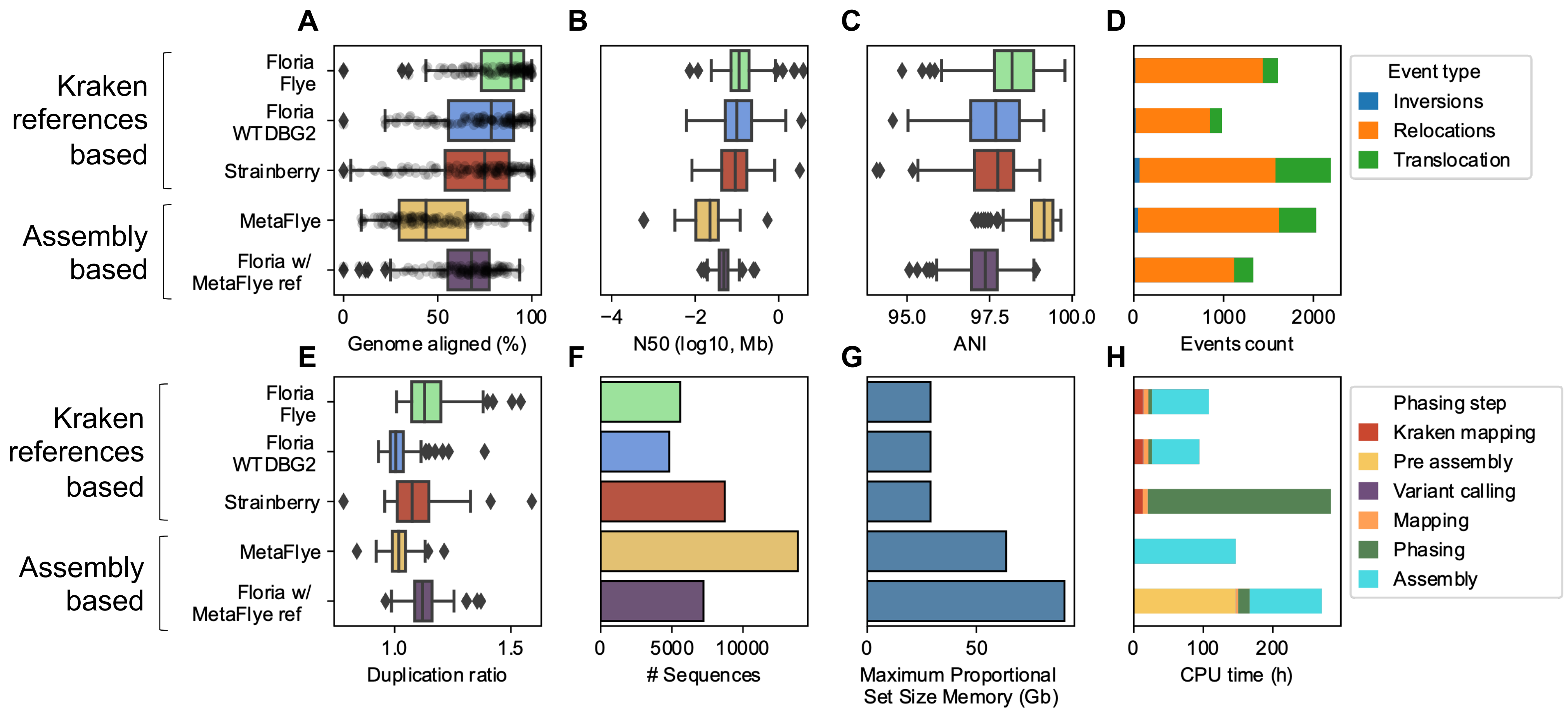


**Supplementary Figure 3. Nanopore phasing assessment results.** Top left to bottom right: (A) Portion of strains genome aligned. (B) N50 of resulting assemblies. (C) ANI of each strain assembly vs the reference sequence of the strain (RSS). (D&E) Number of structural variants observed (D) and duplication ratio (E) of each phased assembly compared to their respective RSS. (F) Number of contigs obtained for each strain assembly. (G) Maximum memory size reached at any point of the pipeline for each phasing methods, adjusted (proportional) to consider shared library. (H) CPU time in hours for each phasing solutions at different steps. For Kraken references based, the assembler below Floria denotes which assembler has been used to assemble Floria haplotigs. Due to its global iterative process, Strainberry is unable to process metagenomic assemblies and is therefore restricted to a references-based approach.

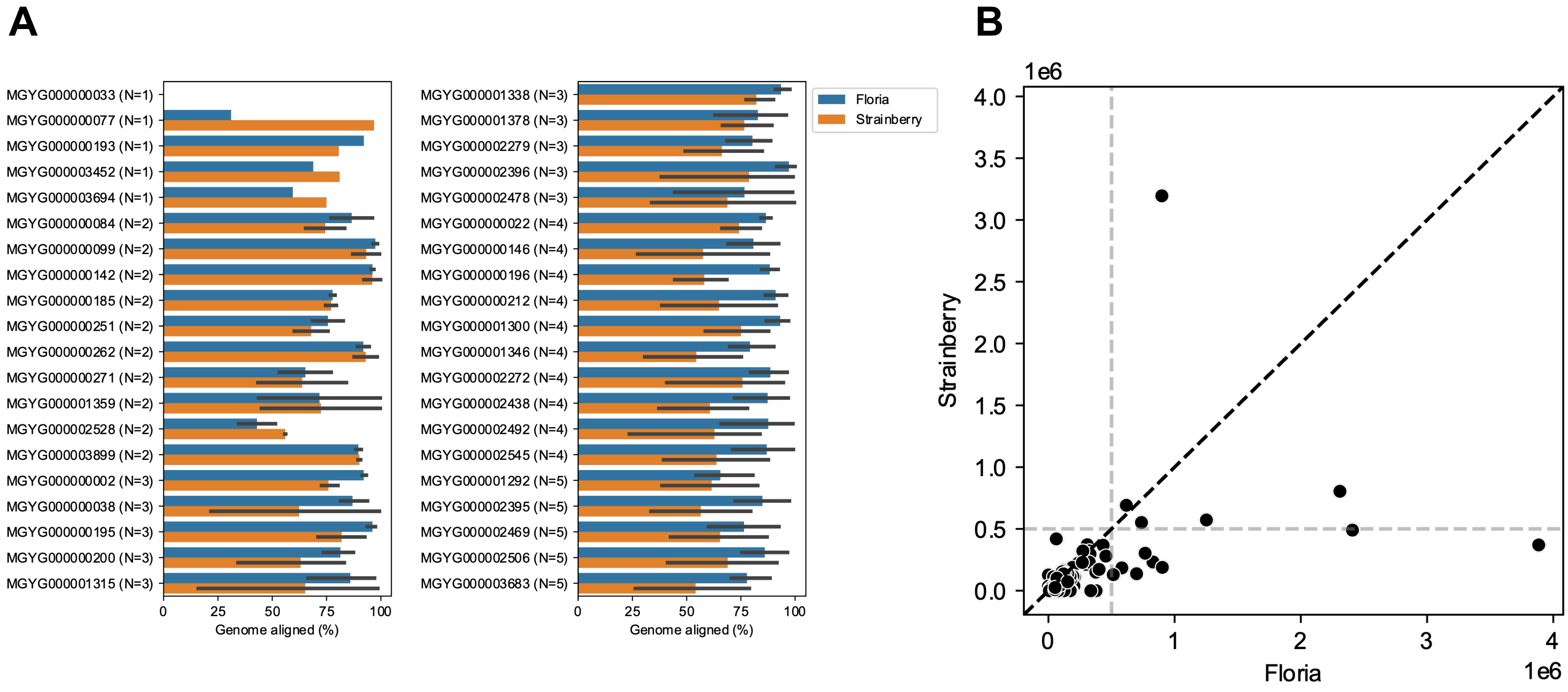


**Supplementary Figure 4. Nanopore phasing comparison.** A. Barplot representation of the percentage of each strain genomes aligned across each processed simulated species. When more than one strain is found within a species, bars represent the mean and additional black line is drawn representing a 95% confidential interval. B. Scatterplot representing the N50 values for both Floria and Strainberry. Grey dashed lines delimit a 500Kb threshold. Black dashed line display graph diagonal.

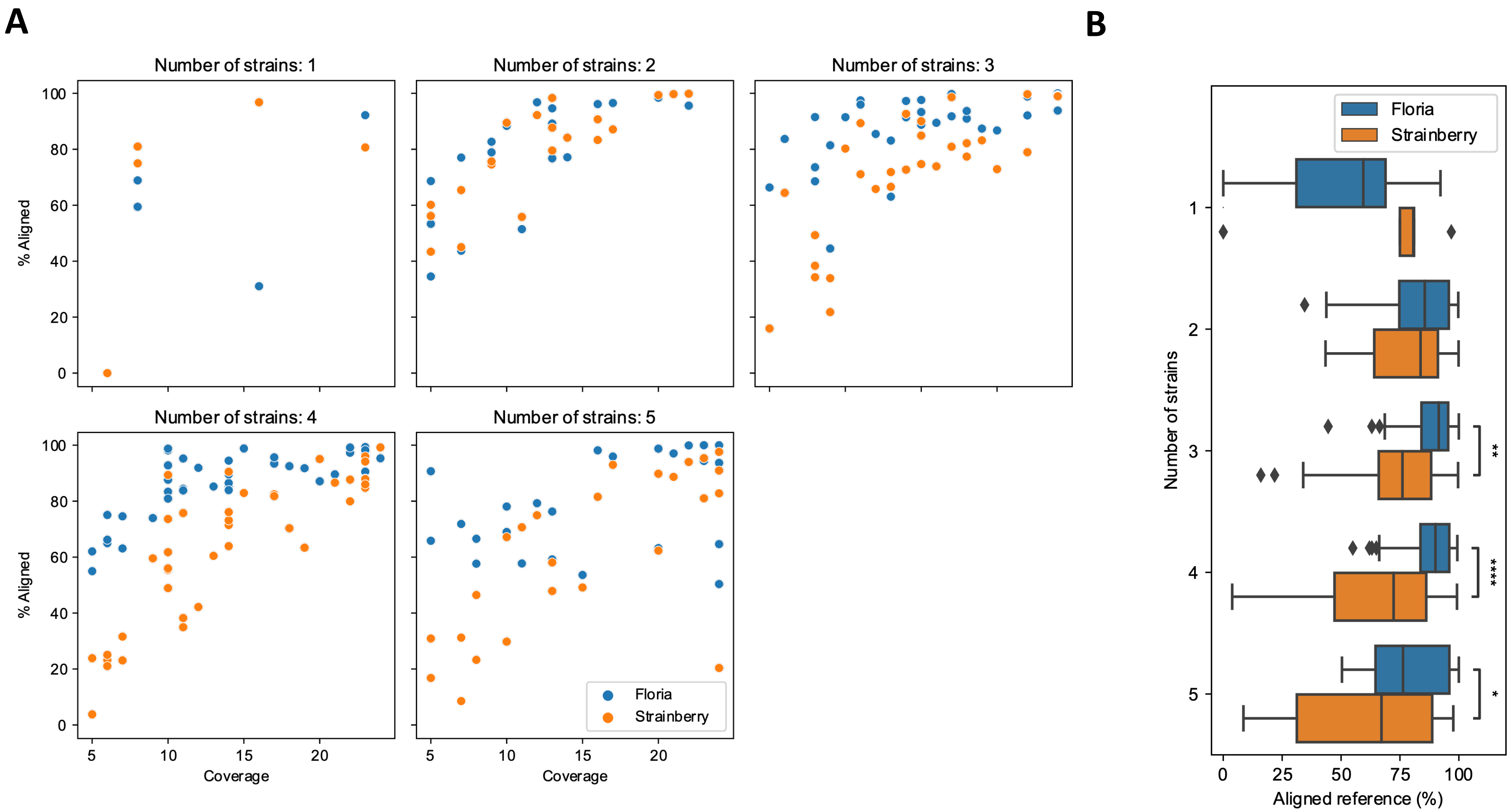


**Supplementary Figure 5. Strains variations within the metagenomic synthetic community.** A. Dot plots representing the percent of aligned reference compared to the strain coverage, grouped by the number of strains used for each species. B. Boxplot representing % aligned reference similar to (A). Significative differences are represented by the * star symbols (Mann-Whitney-U test).

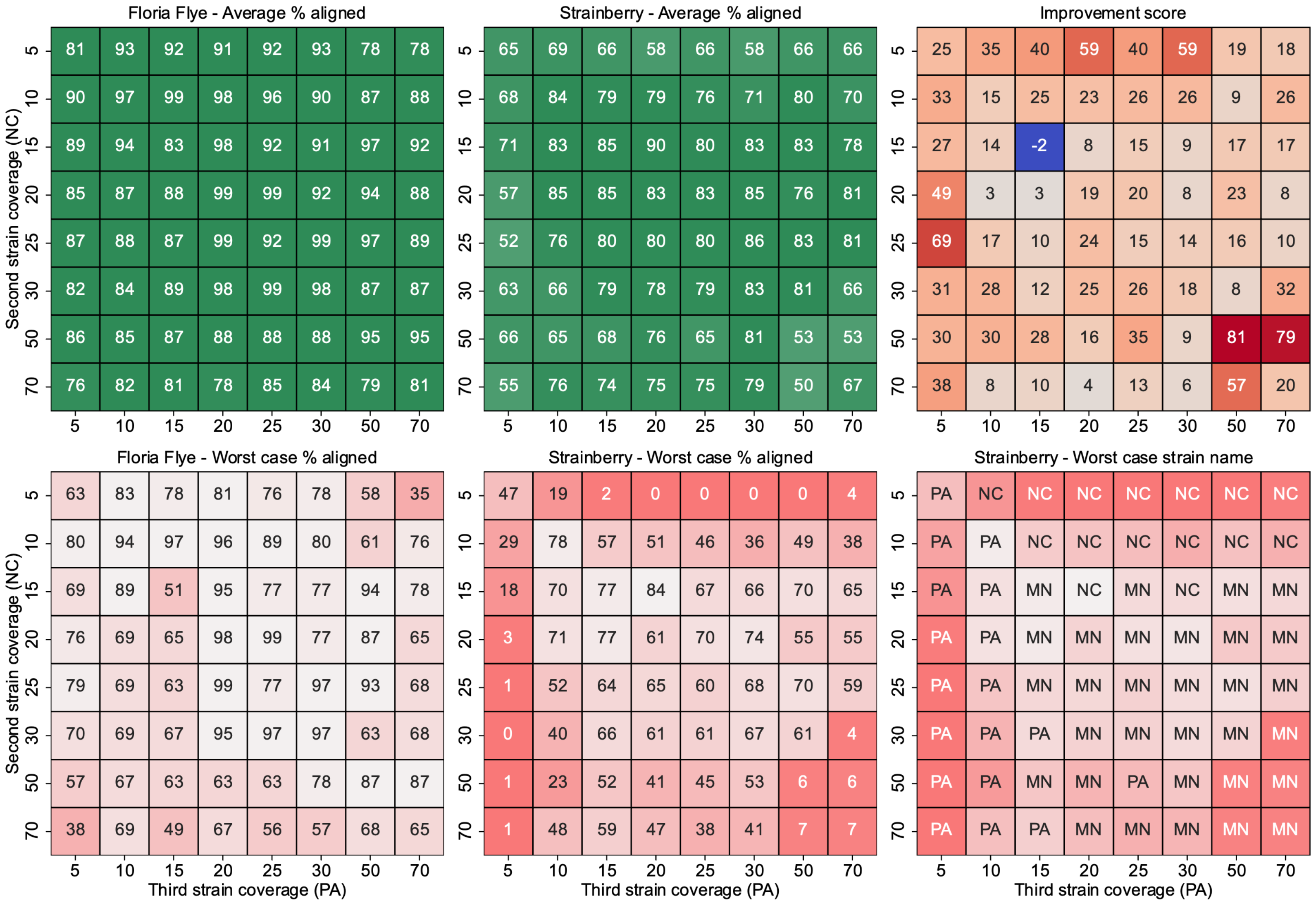


**Supplementary Figure 6. Percent aligned and relative metrics associated for each coverage combination using a *K. pneumonia* three strains community.** Metrics and related softwares are indicated in graph titles. Improvement score is calculated as (floria - strainberry) / strainberry x 100 with percent aligned values. Worst case strain refers to the strains with the worst percent aligned. First strain coverage is 15X.

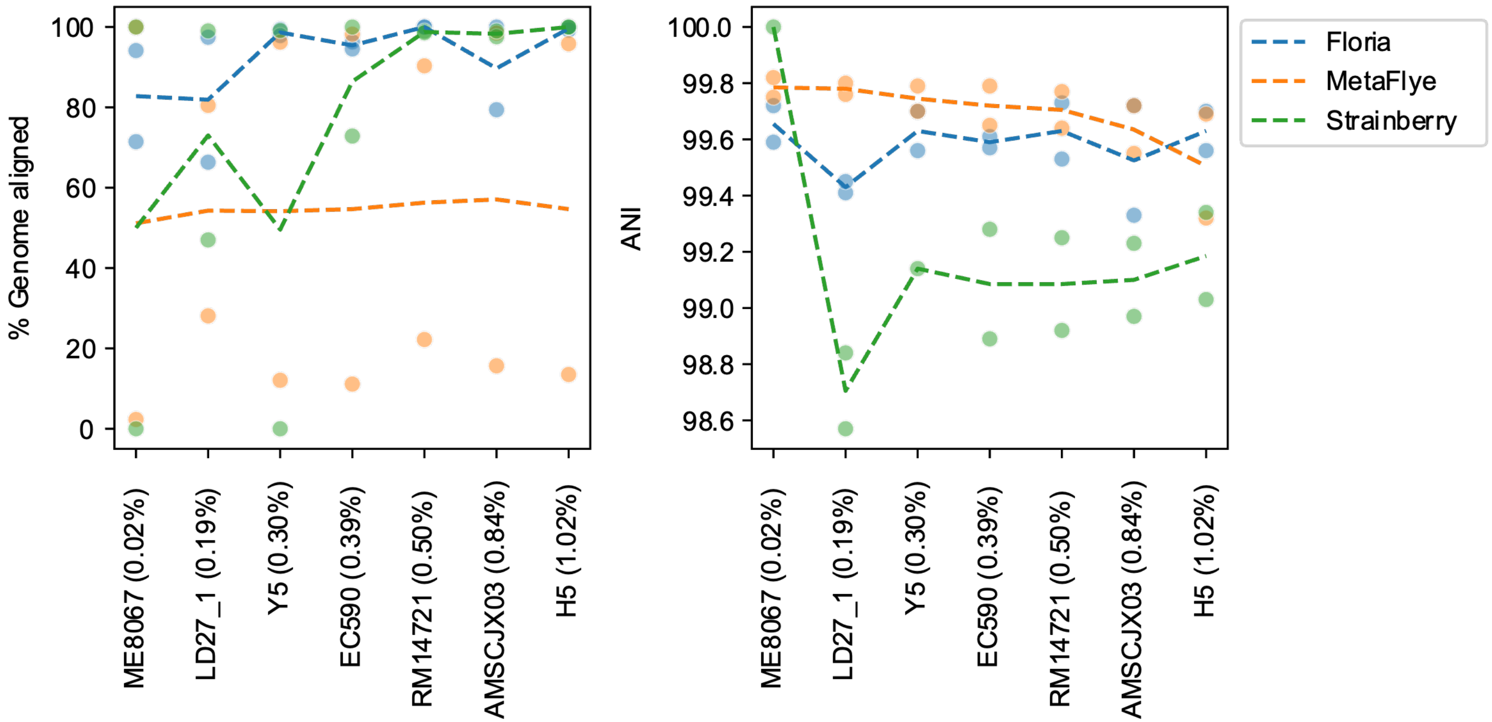


**Supplementary Figure 7. Variable strain divergence.** Percent genome aligned and ANI of multiple two-strains dataset consisting of simulated reads from two strains where one is *E. coli* strain K-12 and the other one is another *E.* coli strain listed on the x-axis (with the divergence percentage shown in parenthesis). Dots represent the exact value of each strain (usually two strains, or one if only one strain genome has been resolved). Dashed lines represent the average value of the one or two strains.

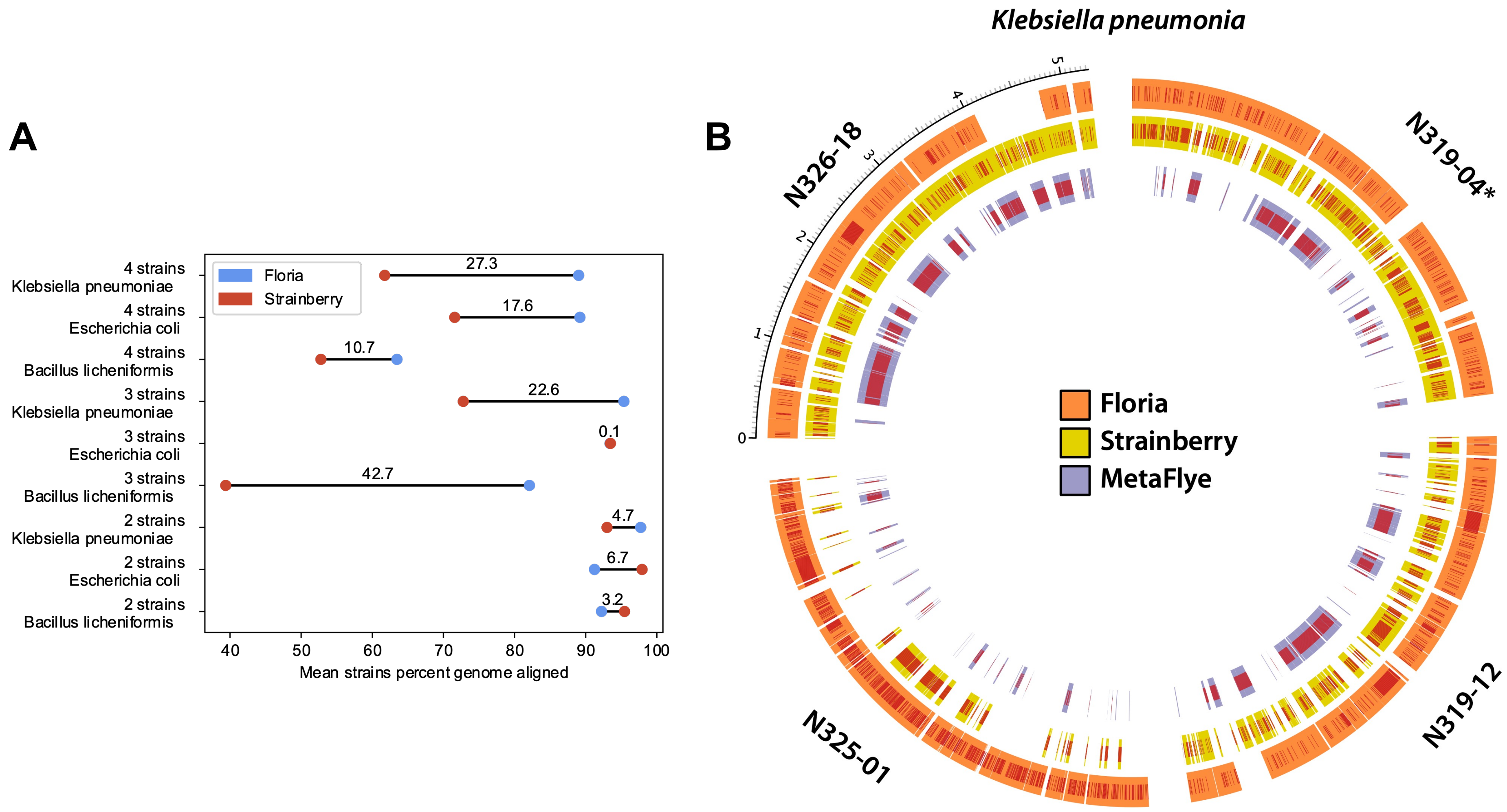


**Supplementary Figure 8. Real reads 3 species communities.** A. Mean strain values of the percent aligned for each species-strains number pair, for both Floria (with Flye) and Strainberry. Numbers between dots represent the absolute difference. B. Circos plot of the *Klebsiella pneumonia* 4 strains community. Bands represent aligned section for each strain genomes and red lines SNPs. Asterisk indicates the reference genome used during the mapping process.

B

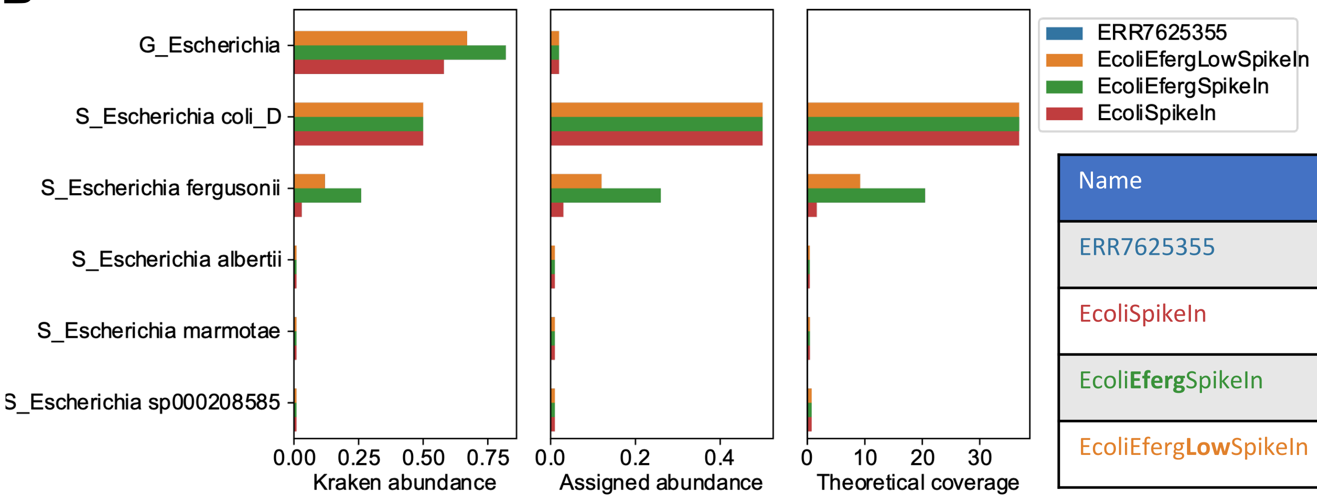


A

| Name | E. coli strain #1 | E. coli strain #2 | E. fergusonii |
|---|---|---|---|
| ERR7625355 | 0 | 0 | 0 |
| EcoliSpikeIn | 20 | 30 | 0 |
| Ecoli**Eferg**SpikeIn | 20 | 30 | 20 |
| Ecoli**Eferg****Low**SpikeIn | 20 | 30 | 8 |

C

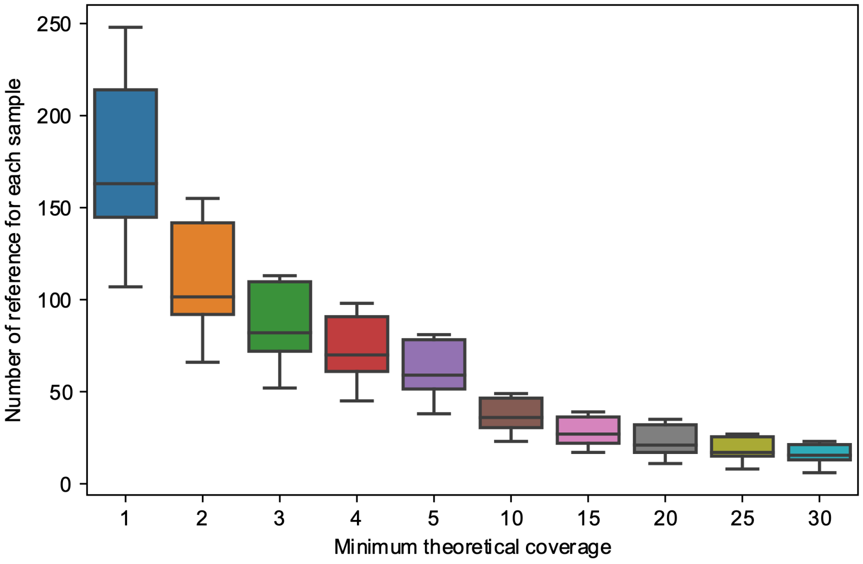


**Supplemental Figure 9. SPMP sample spiked-in results to assess the kraken classification approach.** A. Coverage 4 spike-in datasets. ERR7625355 is the sample used here without spiked-in reads. B: Spike-in results of *E. coli* and *E. fergusoni*. Kraken and assigned abundance, theoretical coverage for each Escherichia species for all 4 datasets described in [A]. C. Number of retained species from the Kraken classification approach based on different minimal theoretical coverage thresholds.

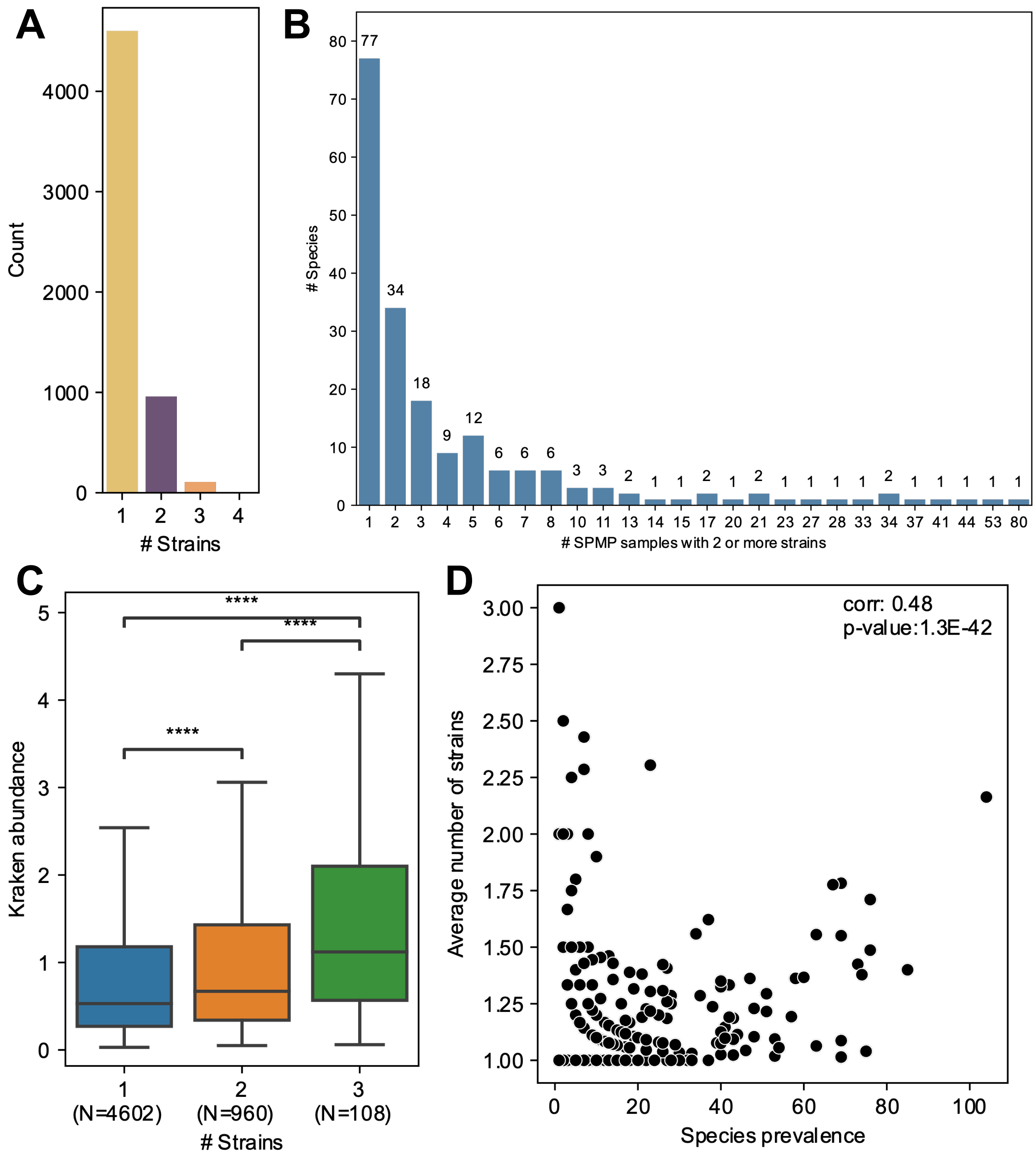


**Supplementary Figure 10. SPMP strains phasing results.** A. Distribution of the number of strains determined within each species. B. Species with more than 1 strain in one sample are more likely to show similar pattern in other samples. C&D. Number of species correlated with species abundance [C] and prevalence [D]. Flyers are ignored in the boxplot representation. Stars (*) represent significative changes (Mann-Whitney-U test). Associated spearman correlation value and p-value [D].

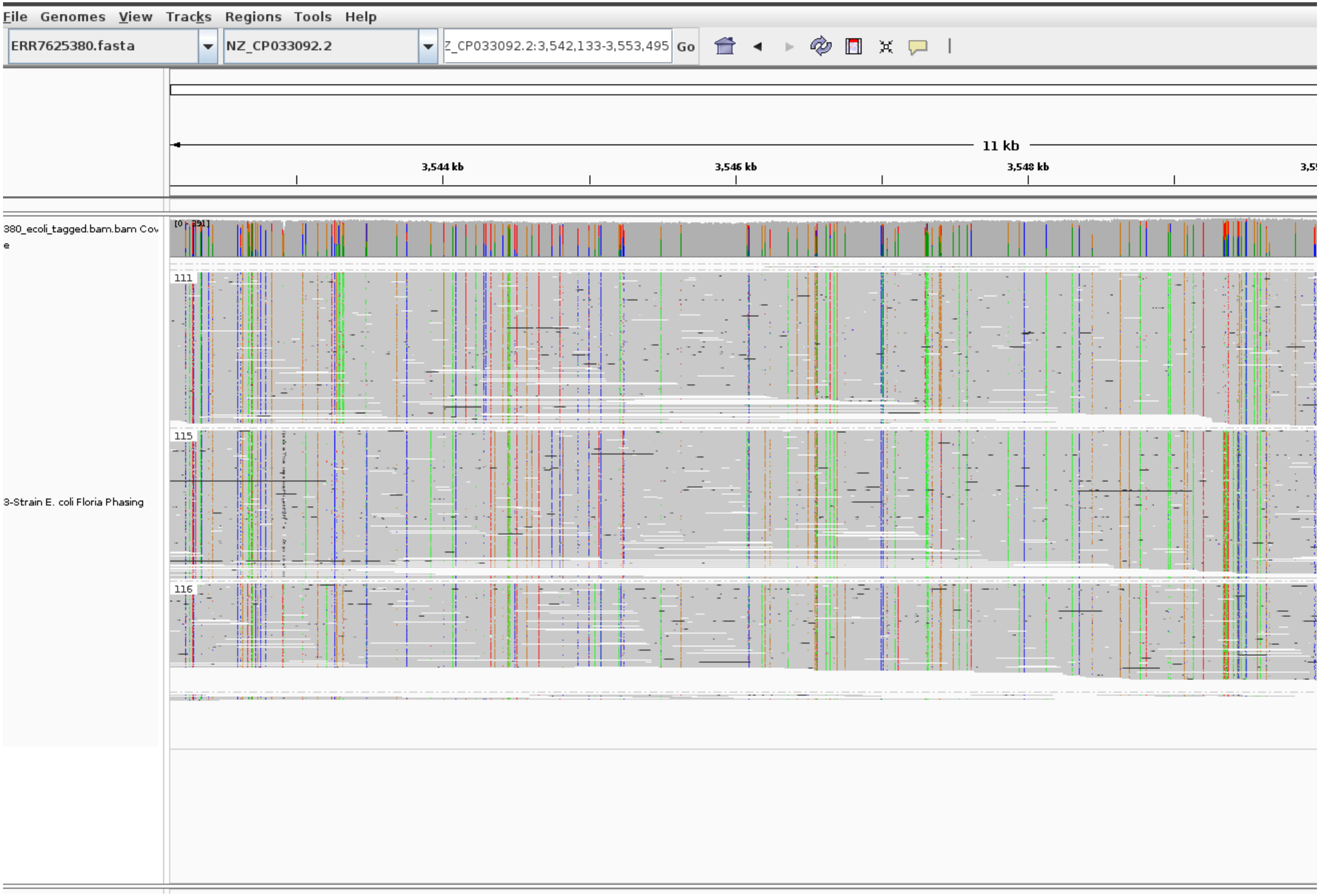


**Supplemental Figure 11**: Haplosets of *E. coli* for sample ERR7625380 from the SPMP cohort. Three haplosets are seen in this section corresponding to three different strains.

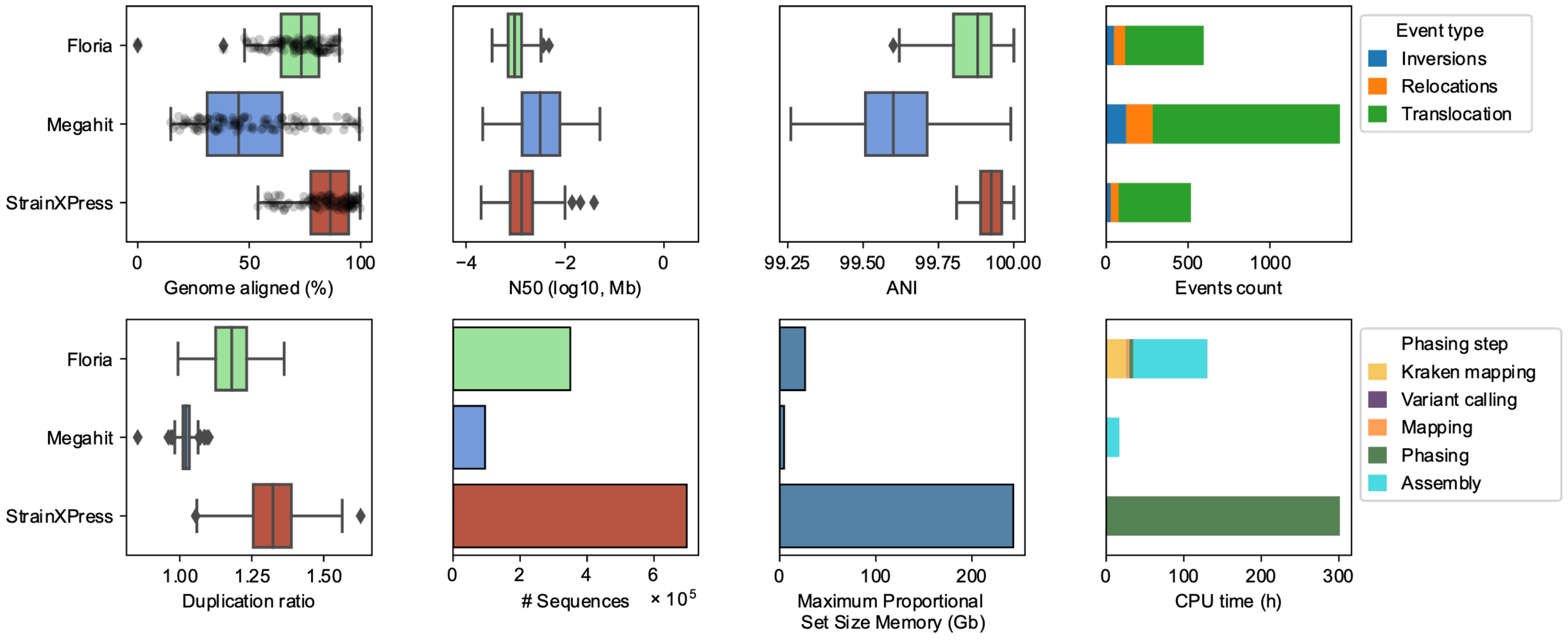


**Supplementary Figure 12. Illumina phasing assessment results.** From top left to bottom right: (A) Portion of strains genome aligned, (B) N50 of resulting assemblies. (C) ANI of each strain assembly vs the reference sequence of the strain (RSS). (D&E) Number of structural variants observed (D) and duplication ratio (E) of each phased assembly compared to their respective RSS. (F) Number of contigs obtained for each strain assembly. (G) Maximum memory size reached at any point of the pipeline for each phasing methods, adjusted (proportional) to consider shared library. (H) CPU time in hours for each phasing solutions at different steps.

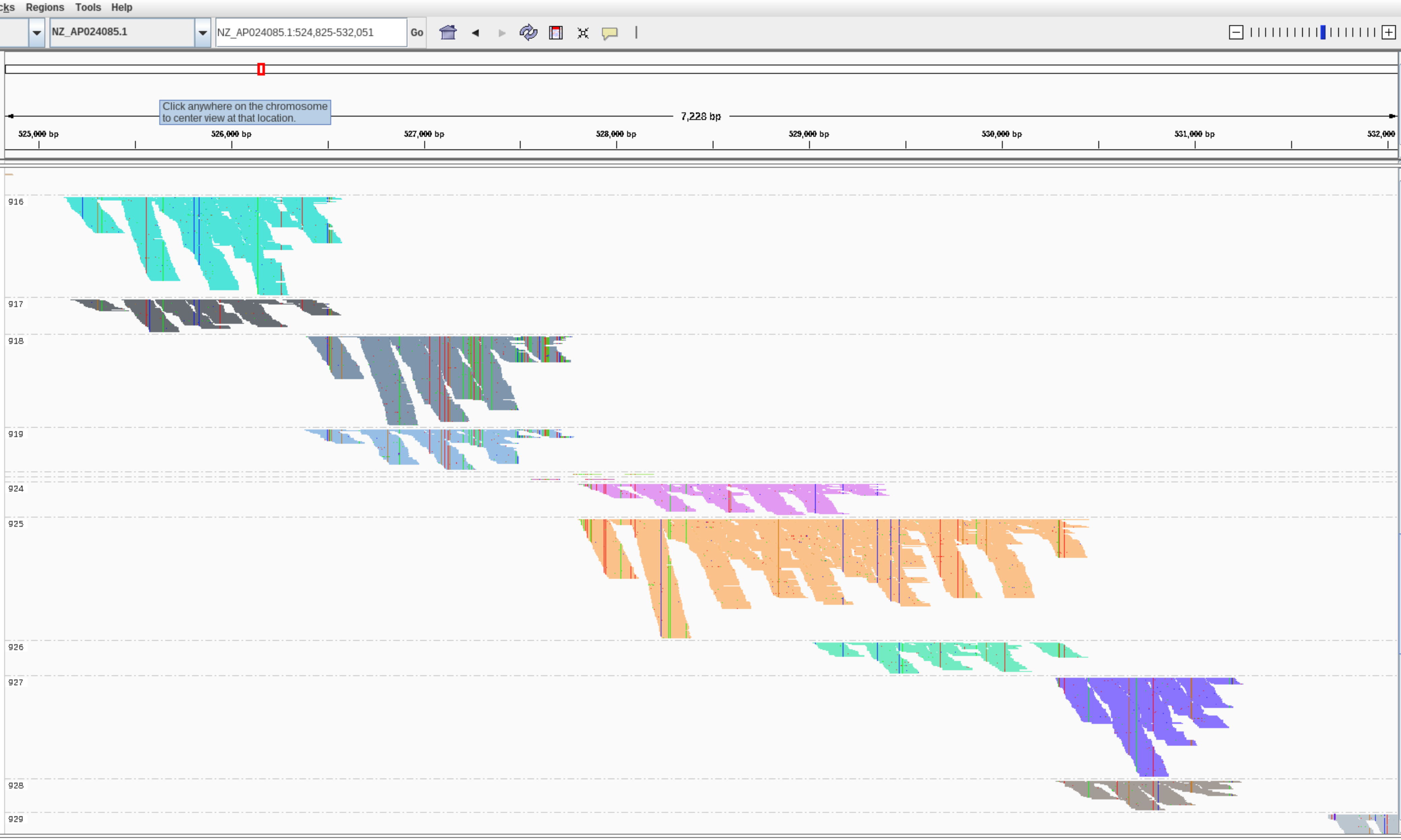


**Supplemental Figure 13**: Haplosets of NZ_AP024085.1, *Faecalibacillus intestinalis*, output by Floria and visualized for the sample dated at day 0 with IGV.

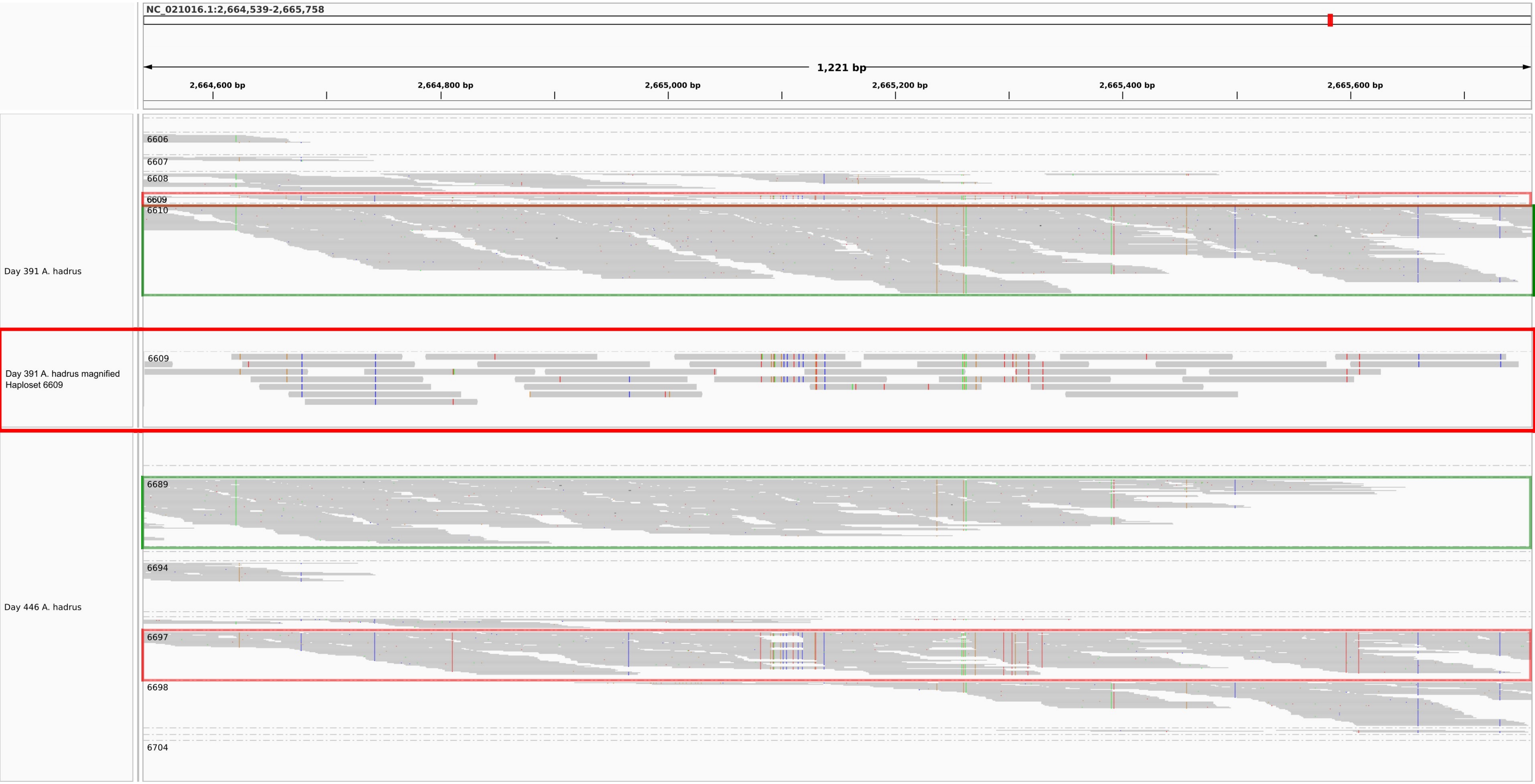


**Supplementary Figure 14**. Haplosets of NC_021016.1, *Anaerostipes hadrus*, output by Floria and visualized with IGV for the sample dated at days 391 and 446. The boxes indicate haplosets that share the same SNPs across timepoints. Note that the red box (haploset 6609) at day 391 is at very low coverage and zooming in confirms that this is the emergent strain from Main Figure 4.

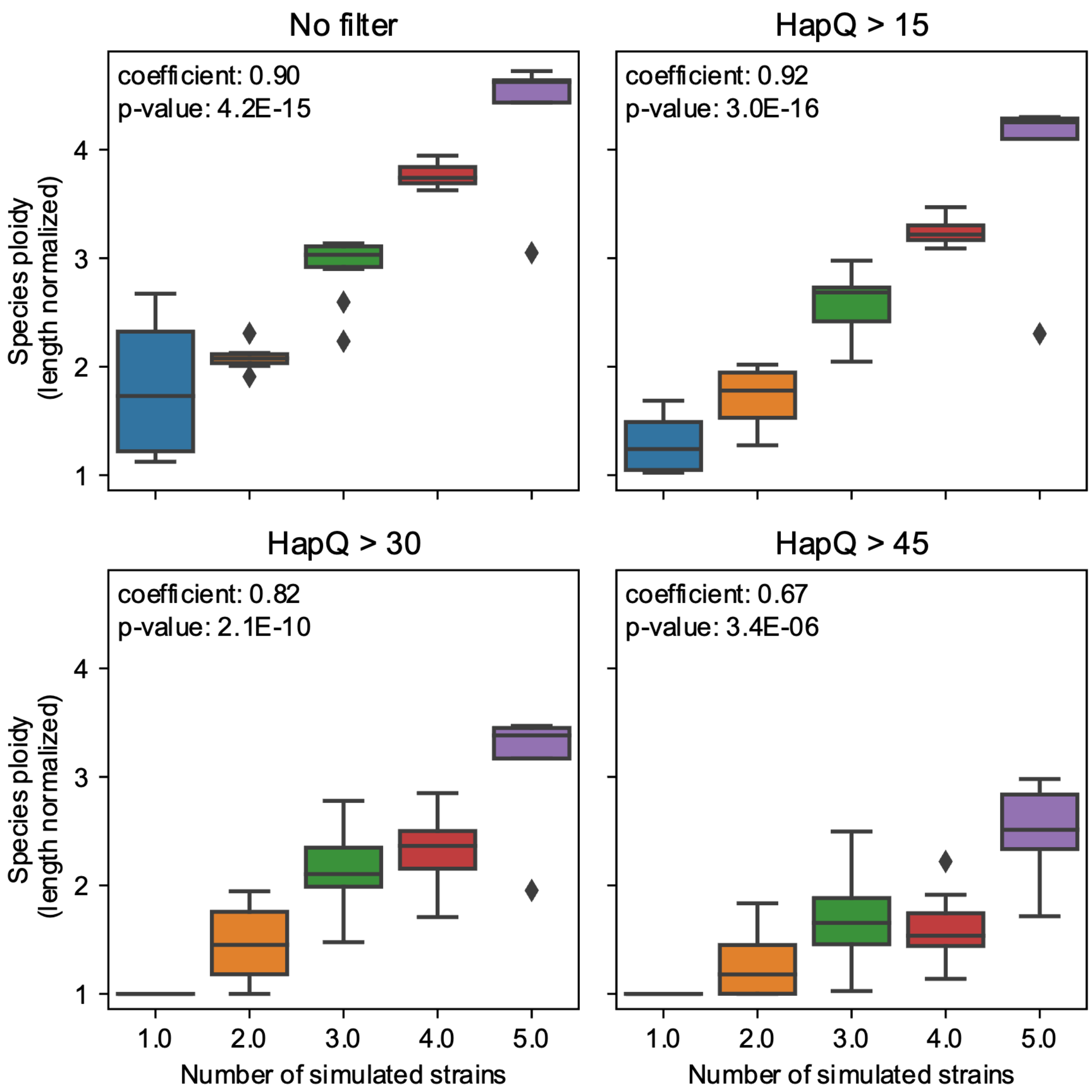


**Supplementary Figure 15**. **Species ploidy estimations of the 40-species synthetic community**. Comparison of the estimated species-ploidy against the number of simulated strains, considering different minimum HAPQ values. Spearman correlation coefficient with associated p-value is annotated on the top left of each plot.
